# Origin of Chromosome 12 Trisomy Surge in Human Induced Pluripotent Stem Cells (iPSCs)

**DOI:** 10.1101/2024.12.02.626470

**Authors:** Maria Narozna, Megan C. Latham, Gary J. Gorbsky

## Abstract

Cultured pluripotent stem cells are unique in being the only fully diploid immortal human cell lines. However, during continued culture, they acquire significant chromosome abnormalities. Chromosome 12 trisomy is the most common whole-chromosome abnormality found during culture of human induced pluripotent stem cells (iPSCs). The conventional paradigm is that trisomy 12 occurs very rarely but provides a proliferative advantage, enabling these cells to outcompete the diploid. Here, we challenge this prevailing model by demonstrating that trisomy 12 arises simultaneously in a very high percentage of diploid cells. Using a single cell line that reproducibly undergoes transition from diploid to trisomy 12, we found that proliferation differences alone do not account for the rapid dominance of trisomic cells. Through careful mapping by fluorescent in-situ hybridization, we identified critical transition passages where trisomic cells first appeared and swiftly gained dominance. Remarkably, single trisomic cells repeatedly emerged de novo from diploid parents. Delving deeper, we discovered an extremely high incidence of chromosome 12 anaphase bridging exclusively during transition passages, along with overrepresentation of chromosome 12 chromatids in micronuclei. These micronuclei fail to replicate during S phase. Subsequently, when these micronucleated cells enter mitosis they contain an unreplicated chromosome 12 chromatids. We also found that nearly 20% of the shorter p arms of chromosome 12 but not the longer q arms exhibited loss of subtelomeric repeats during transition passages. Chromosome 12p arms were exclusively responsible for the bridging observed in anaphase cells. Our findings unveil a novel mechanism of whole-chromosome instability in human stem cells, where chromosome 12p arm-specific segregation errors occur simultaneously in a high percentage of cells. The slight yet significant growth advantage of trisomy 12 cells allows them to persist and eventually dominate the population. Our findings detailing this novel interpretation of the origin of chromosome instability in cultured of human stem cells may have broad implications for understanding the genesis of aneuploidy across diverse biological systems.

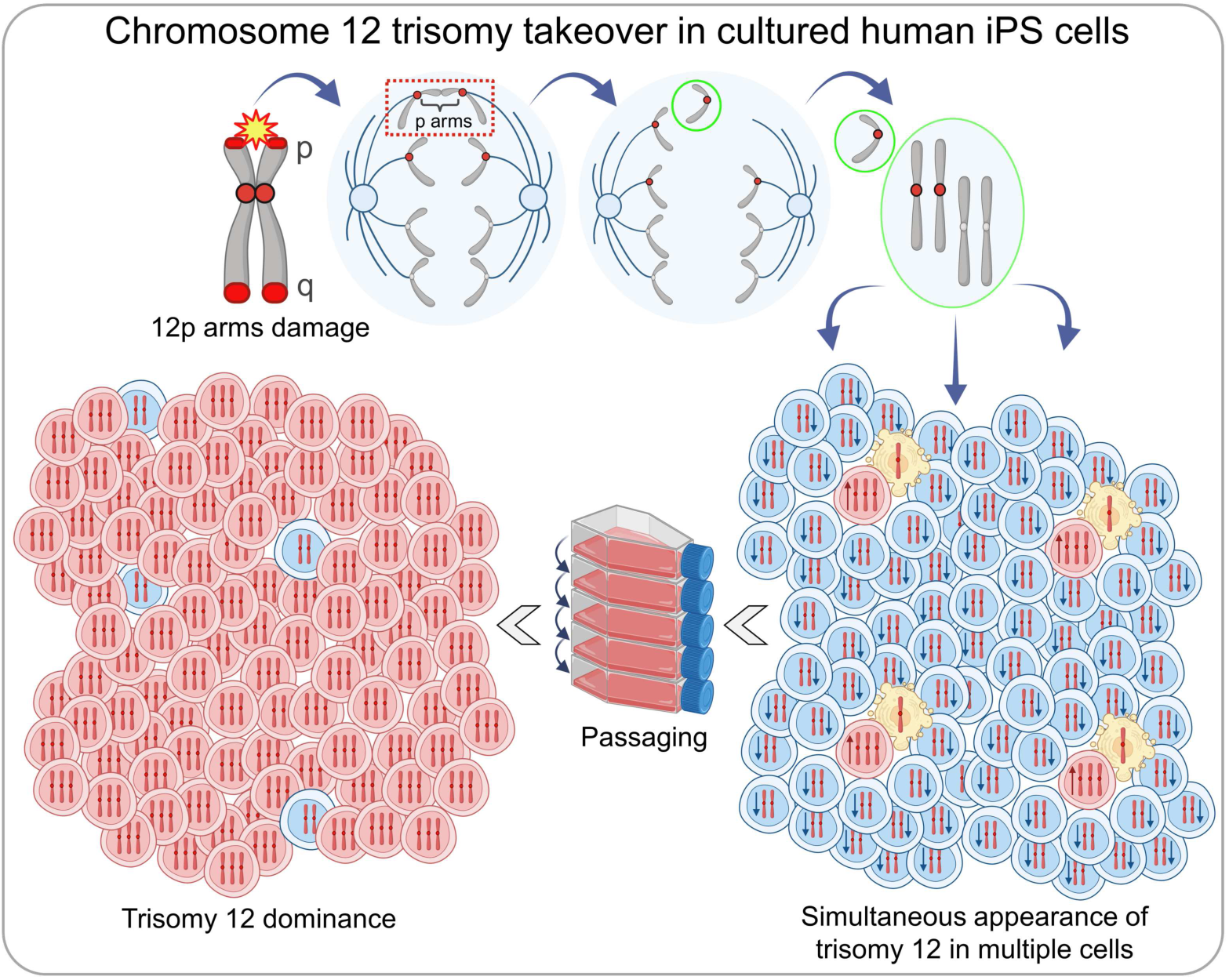

## 1. Introduction

Chromosomal aneuploidy, defined by the gain or loss of whole chromosomes and chromosome segments, is a hallmark of human tumors (*1*, *2*) and presents a major challenge in studying the cell biology of normal diploid cells in cell culture (*3*). Primary cell cultures have limited proliferation potential due to replicative senescence caused by telomere erosion (*4*, *5*). To overcome this, primary cells are immortalized through stable transfection with telomerase (*6*) or with oncogenes (*7*), but the resulting somatic cell lines invariably contain significant chromosome abnormalities (*8-13*). Consequently, so far, most studies of aneuploidy have relied on non-isogenic tumors (*14*) and cell lines (*15*), making it challenging to find isogenic, diploid, and non-transformed human cell line that acquire aneuploidy through natural chromosome mis-segregation to study specific aneuploidies in its native context (*10*).

Human pluripotent stem cells, because they innately express telomerase (*16-18*), are perhaps the only true diploid cells that can be propagated indefinitely in an undifferentiated, pluripotent state in culture (*19*, *20*). Cultured pluripotent stem cells arise from two sources (*21*): embryonic stem cell lines are generated from human embryos (*22*) and induced pluripotent stem cells (iPSCs) are reprogrammed from primary cell cultures via transient expression of specific transcription regulators (*23*). However, up to one-third of all cultured human pluripotent stem cell lines, although initially diploid (*24*), can and do acquire spontaneous chromosome abnormalities over time (*25-29*). These chromosomal abnormalities are not random but predominantly involve duplications of whole or parts of chromosomes 1, 12, 17, 20, and X (*24-31*). Chromosome 12 abnormality most often manifests as a whole-chromosome trisomy, whereas gains involving other chromosomes tend to occur as duplications of single chromosome arms or arm segments (*24*, *27*, *32-34*). The conventional explanation for this phenomenon is random replication and chromosome segregation defects followed by selection of chromosome gains that enhance proliferation (*31-33*, *35-38*). Here we provide evidence that mere slection does not explain the origin and eventual dominance of whole chromosome 12 trisomy in cultured human iPSCs.

In our previous study, we leveraged a single human iPSC line that uniquely acquires trisomy 12 during long-term passaging without other chromosomal aberrations (*39*). Our initial analysis showed that the proliferative advantage of chromosome 12 trisomic cells was insufficient to explain the rapid dominance of the trisomic cells and implicated additional contributors. In our present work, we conclude that the takeover of chromosome 12 trisomy in human iPS cells reflects the gain of an extra copy of chromosome 12 simultaneously in a high percentage of cells. Specifically, we provide a mechanism for the simultaneous conversion of at least 1% of diploid cells to trisomic per each cell division due to a specific chromosome 12-mis-segregation error. Because trisomy 12 cells proliferate faster than their diploid precursors, they are maintained in the population. Eventually, the combination of the generation of new trisomy 12 cells from diploid parents and their faster proliferation rate drive the trisomy 12 cells to rapidly and completely dominate the population. Altogether, our findings uncover a novel source of aneuploidy in human iPSCs with significant implications for the use of stem cells in research and regenerative medicine but may also provide potential intervention strategies to reduce the incidence of spontaneous aneuploidy in stem cell culture.

## 2. Results

### 2.1. Chromosome 12 trisomic cells appear spontaneously in iPS cell culture and rise to almost 100% over 12 3 14 passages

In our previous study, trisomy 12 was first detected at passage 21 in 12% of the cell population and rapidly increased to 80% by passage 26 (*39*). We previously used chromosome spreads to study the shift to trisomy 12, but this approach was limited to small sample sizes. To precisely map the timing of the initial occurrence and eventual dominance of trisomy 12, we used dual-color fluorescence in situ hybridization (FISH) to label centromeric regions of chromosome 12 and, as a control, chromosome 10 (Fig. 1A-C). Chromosome 10 was used as a control due to its comparable size and gene density with chromosome 12 (*40*). Furthermore, karyotypic alterations involving gains or losses of the entire chromosome 10 have not been reported in human iPSCs (*29*, *31*, *41-46*), with only rare instances of 10p loss observed (*25*).

**Fig. 1.**
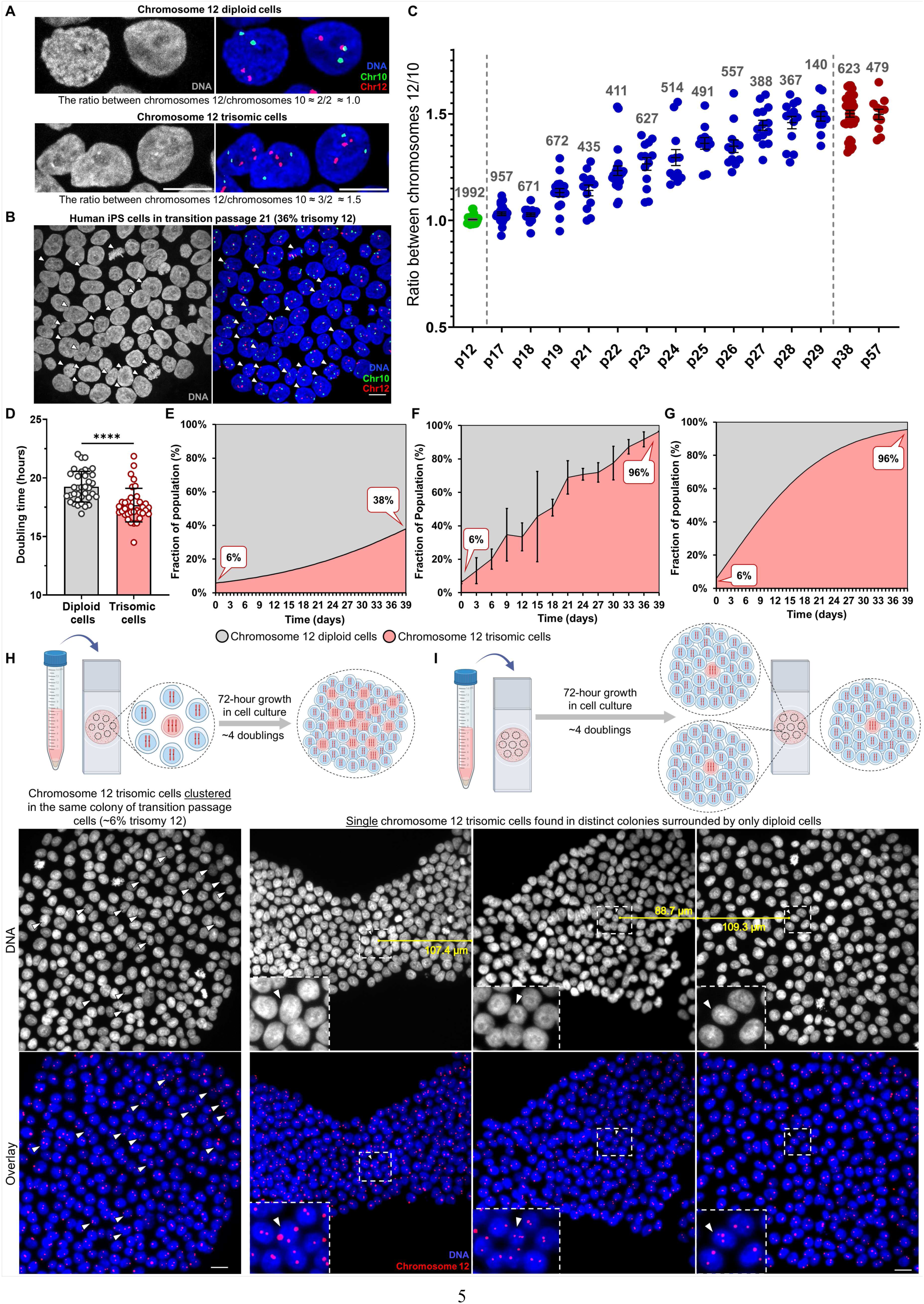
Trisomy 12 dominance in human iPS cells arises spontaneously through multicellular origin and rapidly dominates the cell culture within ∼13 passages. **(A)** Chromosome 12 diploid and trisomic cells after dual-color FISH labeling performed on fixed human iPS cells. DNA labeling is shown in grey. The overlays merge DNA (blue), chromosome 12 centromeres (red), and chromosome 10 centromeres (green). Scale bar = 10 µm. **(B)** Microscope field after FISH performed on fixed human iPS cells from passage 21 with ∼30% of chromosome 12 trisomic cells. Arrows indicate chromosome 12 trisomic cells. DNA labeling is shown in grey. The overlays merge DNA (blue), chromosome 12 centromeres (red), and chromosome 10 centromeres (green). Images were taken using a Zeiss LSM 880 confocal microscope with the Airyscan with 63× objective. Scale bar = 10 µm. **(C)** The observed trisomy 12 takeover during transition passages 17 to 29. Each dot on the graph represents a microscope field of analyzed cells, with the numbers above indicating the total number of analyzed cells. At passage 12, all analyzed cells are still diploid for chromosome 12 (depicted as two red dots labeling two chromosome 12 centromeres) and the control chromosome 10 (two green dots labeling two chromosome 10 centromeres). This results in an average chromosome 12 to chromosome 10 ratio of approximately 1.0. By passage 29 until at least passage 57, nearly all analyzed cells are trisomic for chromosome 12 (possess 3 copies of chromosome 12). This results in a ratio of chromosome 12 to chromosome 10 of 1.5. Transition passages 17-29, show a rapid takeover of the chromosome 12 trisomy. Trisomy 12 persists in the cell culture for at least 28 more post-transition passages since it gained almost 100% dominance at passage 29. Passage numbering reflects the number of passages performed during this study starting from passage 2 of human iPS cell line AICS-0012. Error bars represent SEM. **(D)** Calculated doubling times of diploid and trisomic cell populations (19.25 ± 1.29 h and 17.68 ± 1.41 h, respectively). Data presented as mean ± SD; p values were determined by unpaired two-tailed Mann-Whitney U test, **** p<0.0001. **(E)** Theoretical outgrowth of trisomic cells over diploid during 13 passages (39 days in cell culture) based solely on the doubling times difference. The variable representing the percentage of diploid cells converting to trisomic per each cell division is set here at 0%, relying only on differences in doubling times. **(F)** The averaged dominance of the chromosome 12 trisomic cell population overtime observed in three FISH experiments where trisomy 12 took over the cell culture. Despite expectations of observing 38% trisomy 12 after 13 passages (39 days in cell culture), FISH experiments consistently showed almost 100% trisomic cell population during that time. Error bars represent SEM. **(G)** Trisomy 12 outgrowth modeled by incorporating the doubling time difference and an additional variable representing the percentage of diploid cells converting to trisomic per each cell division. Applying this variable, the mathematical model showed that to reach 96% of trisomy 12 in 13 passages (39 days in cell culture), 3% of diploid cells must convert to trisomic during each cell division. **(H)** Early transition passage (∼6% trisomy 12) showing a predominantly diploid cell colony with clustered trisomic cells, likely arising from a trisomic cell that was seeded among diploids and underwent approximately four doublings over 72 hours of growth. **(I)** Single chromosome 12 trisomic cells observed in various colonies in diploid passage on the same coverslip. In each field, only single trisomic cells were present after three days of growth, with no neighboring trisomic cells detected within at least 90 µm. In **(H)** and **(I)** DNA labeling is shown in grey, with overlays merging DNA (blue) and chromosome 12 centromeres (red). The areas outlined by white boxes were magnified 10x. Images were taken using a Nikon TiE microscope using 40× objective. Scale bars = 20 µm.

It is challenging to find an isogenic cell line in its diploid state and with only trisomy 12 as a sole aberration (*35*). However, the iPS cell line used in our study was analyzed by optical genome mapping and the only large chromosome change detected over long-term passaging was the gain of an additional copy of chromosome 12 (*39*). We used passage 2 of the same, cryopreserved human iPS cell line AICS-0012. We cultured and mapped the cells using FISH over 170 days of continuous culture (57 passages). Cells were passaged every ∼72 hours and grown with Rho-associated protein kinase (ROCK) inhibitor for the first 24 hours post-passaging (*47*). Cell line integrity and authenticity throughout continuous culturing for the FISH experiment were assessed through STR (short tandem repeat) analysis. By analyzing hundreds of cells at each passage, we pinpointed the critical, transition passages where chromosome 12 trisomic cells arose. Chromosome 12 trisomic cells first appeared in a small percentage of cells (∼6%) and gained almost 100% dominance within 12 passages. Trisomic cells remained dominant for at least 28 more post-transition passages (Fig. 1C, S1).

To determine if trisomy 12 detected by centromeric probes reflected the gain of a whole chromosome 12 or a fragment, we used whole chromosome paint for chromosome 12 and applied it to chromosome spreads. This revealed the presence of two complete chromosome 12’s in diploid cells and three in trisomic cells (Fig. S2).

To investigate the robustness and consistency of the trisomy 12 takeover, we conducted independent biological replicates of the previous dual-color FISH mapping, initiated from cryopreserved cells from passage 3 of the AICS-0012 iPS cell line. We continuously passaged the cells every 3 days and mapped nearly every passage for a maximum duration of 45 passages (approximately 135 days). We again identified the critical transition passages where we detected trisomic cells in a small percentage (5-7%) of the cell population (Fig. S3). Trisomic cells progressively dominated the culture over 13-14 passages. The combined findings show that chromosome 12 trisomic cells spontaneously appear in the culture, and once they constitute ∼6% of the population, they rise to nearly 100% within ∼13 passages.

Overall, we have cultured the same AICS-0012 iPS cell line through transition passages a total of eight times, once in the previously published study (*39*) and seven times more recently. In seven out of eight experiments, trisomy 12 became dominant to over 95% of the population, demonstrating that this is the most frequent outcome. In a single replicate, we did not observe significant trisomy 12 (Fig. S4A-B), and we considered the possibility that the cell population acquired other major chromosomal aberrations during culturing. However, KaryoStat+ analysis revealed no additional chromosomal gains or losses indicating that in rare cases, iPSC cultures do not acquire significant chromosomal aberrations (Fig. S4C).

### 2.2. The difference in doubling times between chromosome 12 diploid and trisomic cells of the same origin is insufficient to account for the rapid dominance of the trisomic population

In our previous study, the change in growth rate due to amplification of chromosome 12 in cell culture was insufficient to explain the dominance of trisomic cells. However, the growth rates of chromosome 12 diploid and trisomic cells calculated in that study were based only on cell counting during passaging (*39*). To obtain high-resolution proliferation measurements, we performed microwell growth assays using SYBR Gold (*48*, *49*) and determined doubling times for the diploid and trisomic populations (Fig. S5A-C). We found that diploid cells showed a doubling time of 19.25 ± 1.29 hours (mean ± S.D.), while trisomic cells 17.68 ± 1.41 hours (Fig. 1D). These results are consistent with the literature data indicating human iPS cells division time of 18-20 hours (*39*, *50*).

We used doubling times to model the theoretical outgrowth of the trisomic population over diploid, starting from an initial average 6% trisomic cells in the first transition passage. Based on the doubling times, we would have expected only 38% of trisomic cells after 13 passages (39 days in the cell culture) (Fig. 1E). However, our experiments consistently showed almost 100% trisomy 12 at that time (Fig. 1F), confirming that differences in doubling times alone cannot explain the rapid takeover by trisomic cells. We then introduced to the model a variable representing the percentage of diploid cells converting to trisomic per cell division. Our modeling showed that to reach 96% of trisomy 12 in 13 passages (as observed in FISH experiments), 3% of diploid cells must convert to trisomic during each population doubling (Fig. 1G).

To verify whether trisomy 12 cells arise simultaneously in multiple cells, we examined diploid and early transition passages. In early transition passages (∼6% trisomy 12), we confirmed that primarily diploid cell colonies contained clusters of trisomic cells (Fig. 1H). This clustering suggests that trisomic were seeded and grew for 3 days, dividing approximately four times to form colonies with grouped daughter trisomic cells. However, after 3 days of growth, we also found multiple, isolated, single trisomic cells (Fig. 1I). These trisomic cells must have arisen less than one doubling time before fixation. Altogether, our data indicate that trisomy 12 arises simultaneously and independently in multiple cells.

### 2.3. Chromosome 12 shows a high propensity for bridging during anaphase in transition passage iPSCs

To understand how chromosome 12 trisomy arises in transition passages we examined the segregation of chromosome 12 and control chromosome 10 in mitosis. Exclusively during the transition passages, 55% of detected anaphase bridges involved chromosome 12, a frequency 13-fold higher than expected by random chance (approximately 4.3% for any of the 23 human chromosomes). This level of bridging was never observed for chromosome 10. However, once trisomy 12 became dominant (in ∼100% trisomic passages), the tendency for chromosome 12 to form bridges significantly decreased. In very early fully diploid passages, chromosome 12 bridging was not elevated (Fig. 2A-B).

**Fig. 2.**
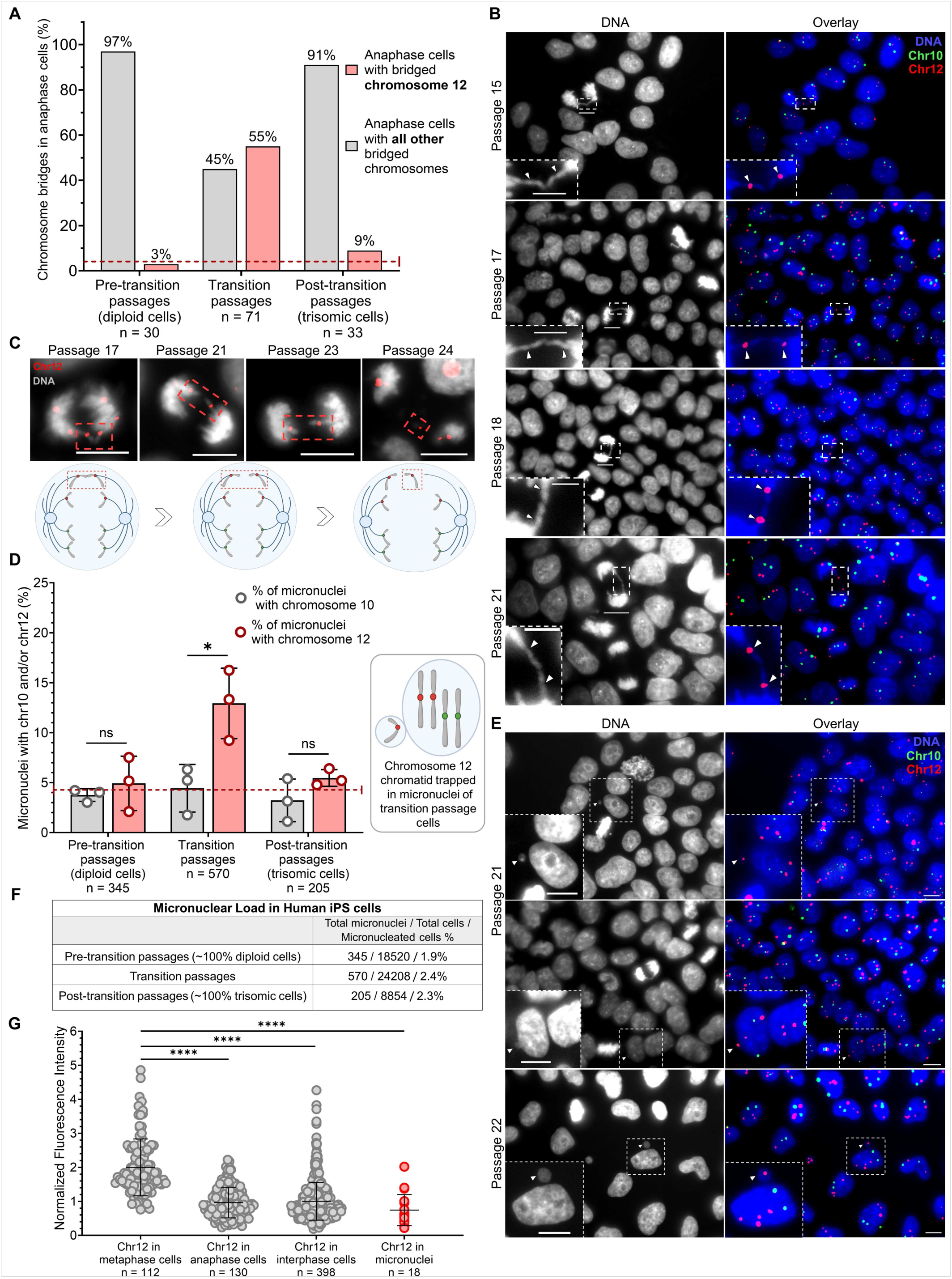
During transition passages only, chromosome 12 shows a high propensity for bridging during anaphase and is significantly overrepresented in micronuclei. **(A)** The percentage of bridged chromosome 12 in pre-transition (∼100% diploid), transition, and post-transition (∼100% trisomic) iPS passages. In transition passages, bridging chromosome 12 exceeds to the highest extent the theoretically assumed percentage of each chromosome bridging in anaphase (1 out of human 23 chromosomes randomly bridging would give approximately a 4.3% chance). The n values represent the number of analyzed bridges. The dotted line indicates the random expected frequency for any single chromosome bridging in anaphase. **(B)** Anaphase cells with one copy of chromosome 12 bridging found in transition passages 15, 17, 18, and 21. DNA labeling is shown in grey, the overlays merge DNA (blue), chromosome 10 centromeres (green), and chromosome 12 centromeres (red). Scale bars = 10 µm. Scale bars in zoomed regions = 5 µm. **(C)** Delayed separation of chromosome 12 chromatids observed in transition passages 21, 23, and 24 anaphase cells, possibly due to prior bridging. The overlays merge a single plane of DNA labeling (gray) with chromosome 12 centromeres (red). Red-boxed areas indicate the bridged and lagging chromatids. Scale bars = 10 µm. **(D)** The percentage content of chromosome 10 and chromosome 12 in micronuclei found in diploid cells (pre-transition passages), transition passage cells, and trisomic cells (post-transition passages) in three replicates of FISH trisomy 12 takeover experiments. The dotted line indicates the random expected frequency for any single chromosome being mis-segregated into the micronuclei (1 chromosome out of 23 = 4.3%). The percentage of chromosome 12 in micronuclei of transition passages exceeds the randomly expected percentage of 4.3% to the greatest extent. The n values represent the number of analyzed micronuclei in three replicates of FISH trisomy 12 takeover experiments. Data presented as mean ± SD; p values were determined by unpaired two-tailed Student’s t-test with Welch correction, * p=0.0318. **(E)** Representative micronuclei observed in transition passages 21 and 22 containing chromosome 12 chromatids. In FISH, identifying cells in S or G2 phases - where chromosome 10 and/or 12 are replicated and exhibit distinct signals - is very rare. However, passage 22 depicts chromosome 12 diploid cell with already replicated chromosome 12 (4 centromeric signals). Remarkably, the fifth chromosome 12 signal of comparable fluorescence intensity to those in the main nucleus originates from the micronucleus, suggesting the presence of a fifth chromosome 12 chromatid in the cell. DNA labeling is shown in grey, the overlays merge DNA (blue), chromosome 10 centromeres (green), and chromosome 12 centromeres (red). Scale bars = 10 µm. **(F)** Counts of micronucleated cells as a fraction of the total number of micronuclei and the total number of analyzed cells in experiments when trisomy 12 took over the culture. The values for pre-transition passages (18520 cells), transition passages (24208 cells), and post-transition passages (8854 cells) represent the cumulative results of three independent counts from three separate FISH experiments. **(G)** Fluorescence intensity (F.I.) of chromosome 12 centromeres in metaphase cells was normalized to 2. Each dot in the metaphase group (n = 112) represents the F.I. of two adjoined chromosome 12 centromeres. Chromosome 12 F.I. in anaphase cells (n = 130), as well as in interphases (n = 398) and micronuclei (n = 18) was normalized to metaphase cells. Each dot in these groups represents the F.I. of an analyzed signal. Data presented as mean ± SD; p values were determined by unpaired two-tailed Mann-Whitney U test, **** p<0.0001.

An anaphase chromosome bridge may occur when cells enter mitosis with fused sister chromatids resulting in lagging of the bridged chromosomes in anaphase. This lag can position bridged chromosomes near the spindle equator, leading to mis-segregation of both chromatids into the same daughter nucleus or into a micronucleus (*51*). In cells at late anaphase, we also detected separated but lagging chromosome 12 chromatids consistent with the idea that the bridged chromatids eventually rupture their connection and separate (Fig. 2C).

### 2.4. Chromosome 12 is entrapped in micronuclei at high frequency

We reasoned that bridging chromosome 12 chromatids can selectively become entrapped in micronuclei of transition passage cells. Micronuclei are widely recognized and studied as biomarkers of chromosome instability observed in cancer, with most arising from chromosome mis-segregation due to mitotic errors (*52*). While micronuclei have been extensively studied in cancer cells (*53*), they are less explored in human pluripotent stem cells. The high proliferation rates of human pluripotent stem cells make them particularly susceptible to micronuclei formation (*42*, *54*).

We compared the rates of mis-segregation of chromosomes 10 and 12 into micronuclei during continuous culture (Fig. 2D-E). We found that chromosome 12 was enriched in micronuclei only in transition passages, where its mis-segregation rate (12.9%, p=0.0318) was significantly higher than that of chromosome 10 (4.4%). In contrast, in pre-transition and post-transition passages, the mis-segregation rates for chromosomes 10 and 12 were similar and not significantly different (Fig. 2D). Micronuclei were consistently observed across pre-transition, transition, and post-transition cells, with no significant change in their frequency (Fig. 2F).

To investigate whether chromosome 12 signals in the micronuclei reflected single chromatids or whole chromosomes we devised a method for counting chromatids using quantitative fluorescence. We generated standards by measuring the fluorescence signals exhibited on metaphase chromosomes that contain two centromeres and signals from anaphase chromosomes that contain single centromeres. Our analysis showed that the fluorescence of chromosome 12 centromere in micronuclei was significantly lower (p<0.0001) than the intensity of metaphase chromosomes and comparable to the intensity of anaphase chromatids (Fig. 2G), indicating that these signals originate from single chromatids.

### 2.5. The presence of intact lamin B1, γ-H2AX expression, and the percentage of large micronuclei indicate that over 50% of micronuclei in human iPSCs remain stable

Micronuclei are a hallmark of chromosome instability in cancer cells where they often exhibit defects in nuclear lamina, leading to rupture of their envelopes, followed by enzymatic shattering of the enclosed chromosomes, a process termed chromothripsis (*53*, *55-60*). We examined the distribution of lamin B1 to determine nuclear lamina integrity in micronuclei of human iPSCs. We used diploid and early transition passages of the AICS-0013 cell line, which carries an mEGFP tag on the N-terminus of the lamin B1 gene *LMNB1*. We observed that lamin B1 was present in all primary nuclei and approximately 60% of micronuclei (Fig. 3A-C). Given that the frequency of micronuclei rupture correlates with the presence of lamina gaps (*61*), we analyzed 3D projections of micronuclei. Micronuclei exhibiting gaps in the nuclear lamina meshwork, which create weak points susceptible to membrane rupture (*56*, *61*), were scored as lamin B1-negative.

**Fig. 3.**
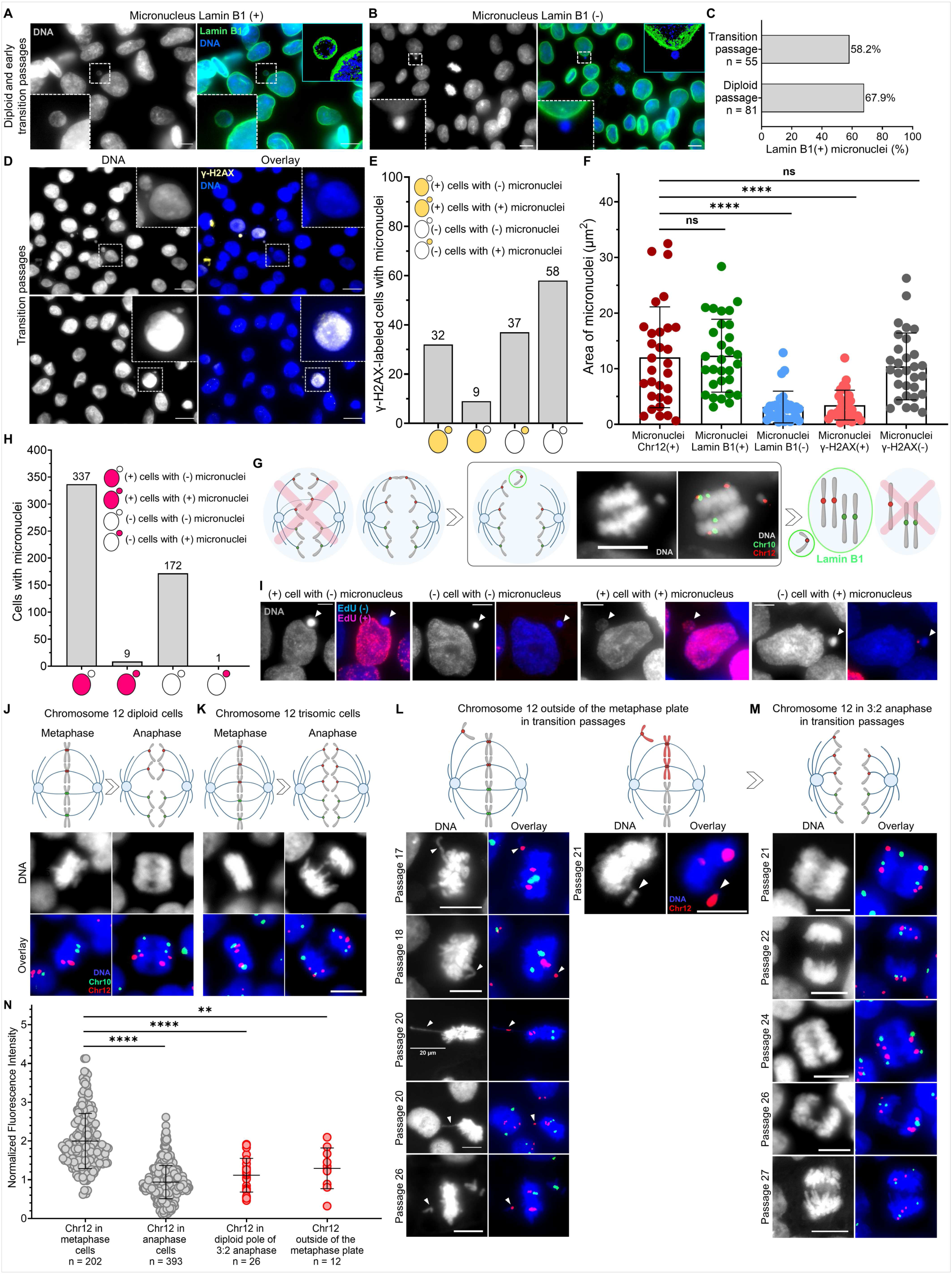
Transition passage cells enter mitosis with one unpaired chromosome 12 chromatid originating from prior mis-segregation into micronuclei. **(A)** and **(B)** Representative human iPS cells micronuclei with and without lamin B1 expression in the micronuclear membrane. 3D projections, generated using MetaMorph software were used to assess lamina integrity. DNA is shown in grey, and the overlay represents DNA (blue) and lamin B1 (green). The white box areas were magnified 10x. Scale bars = 10 µm. **(C)** Percentage of lamin B1 presence in micronuclei of diploid and early transition passages. GFP visualization of the LMNB1-GFP-tagged iPS cell line showed that lamin B1 was present in all primary nuclei and approximately 60% of micronuclei. The n values represent the number of analyzed micronuclei. **(D)** Representative images of γ-H2AX immunolabeling of transition passage iPSCs. DNA labeling is shown in grey, and the overlay represents DNA (blue) and γ-H2AX (yellow). The white box areas were magnified 10x. Scale bars = 10 µm. **(E)** Human iPS cells with micronuclei in transition passages analyzed for γ-H2AX presence in both primary nuclei and micronuclei. Of over 130 cells with micronuclei, approximately 50% had γ-H2AX-negative micronuclei, the majority of which were also negative for the presence of double-strand breaks in the main nuclei. **(F)** Area of micronuclei measured across five groups, with the first group representing micronuclei containing mis-segregated chromosome 12 chromatids. Micronuclei with chromosome 12 were compared to lamin B1-positive, lamin B1-negative, γ-H2AX-positive, and γ-H2AX-negative micronuclei. Micronuclei containing chromosome 12 were significantly larger than lamin B1-negative and γ-H2AX-positive micronuclei but were not significantly different from lamin B1-positive or γ-H2AX-negative micronuclei. Data presented as mean ± SD (n = 30 for each group); p values were determined by unpaired two-tailed Mann-Whitney U test and Student’s t-test with Welch correction, **** p<0.0001. **(G)** Model of chromosome mis-segregation and lamin B1 recruitment in the subsequent interphase, adapted from Liu et al. (*59*). Micronuclei formed outside the spindle are more stable, as they recruit non-core proteins, such as lamin B1. Lagging chromosome 12 was never observed within the midspindle. However, when chromosome 12 bridges or mis-segregation events were observed, they predominantly occurred as peripheral bridges or mis-segregation outside the midspindle, as illustrated in the example anaphase cell. DNA labeling is shown in grey and the overlay merges DNA (grey), chromosome 12 centromeres (red), and chromosome 10 centromeres (green). Scale bar = 10 µm. **(H)** Human iPS cells with micronuclei in transition passages with incorporation of EdU showing DNA replication in nuclei and micronuclei within the same cells. The most frequent outcome was EdU incorporation into primary nuclei but not micronuclei within the same cells. **(I)** The four scored patterns of EdU incorporation in iPSCs with micronuclei after pulse labeling with 20 μM EdU for 20 min. DNA labeling is shown in grey, the overlay merges DNA (blue) and EdU labeling (red). Scale bars = 5 µm. **(J)** and **(K)** show schematics and the corresponding FISH labeling on the regular metaphase and anaphase cells of chromosome 12 diploid and trisomic cells. **(L)** Metaphase cells observed in transition passages with signals from one potentially unpaired chromosome 12 chromatid apart from the metaphase plates. In metaphase cells, two copies of chromosome 12 are found in the aligned metaphase plate while the third copy lies at some distance off the metaphase plate. In a separate experiment using whole chromosome 12 probes, again two copies of chromosome 12 are found in the aligned metaphase plate (one is weakly visible due to being positioned deeper within the metaphase plate), while the third copy lies off the metaphase plate. **(M)** Anaphase cells from multiple transition passages with the normal 2 and 2 distribution of control chromosome 10 chromatids, but anomalous organization of 3 chromatids of chromosome 12 oriented to one pole and 2 chromatids oriented to the other pole (3:2 anaphase cells). In **(D)**-**(G)**, DNA labeling is shown in grey and the overlays merge DNA (blue) and centromeric probes for chromosome 10 (green) and chromosome 12 (red). The right overlay in panel F overlays DNA (blue) and whole chromosome paint for chromosome 12 (red). Scale bars = 10 µm. **(N)** Fluorescence intensity (F.I.) of chromosome 12 centromeres in diploid and trisomic metaphase cells was normalized to 2. Each dot in the metaphase group (n = 202) represents the F.I. of two adjoined chromosome 12 centromeres. Chromosome 12 F.I. in diploid and trisomic anaphase cells (n = 393), as well as in diploid poles of 3:2 anaphase cells (n = 26) and outside of the metaphase plates (n = 12) was normalized to metaphase cells. Each dot in these groups represents the F.I. of an analyzed signal. Data presented as mean ± SD; p values were determined by unpaired two-tailed Mann-Whitney U test. P-values are as follows: ** p=0.0017, **** p<0.0001.

Smaller micronuclei have been reported to inefficiently form intact lamina, making them prone to rupture and accumulation of DNA damage (*62*). We found that the presence of an intact lamin B1 shell correlates with larger micronuclei size. The average area of lamin B1-positive micronuclei was 12.34 ± 6.39 μm^2^ (mean ± S.D.) and the average area of lamin B1-negative micronuclei was 3.12 ± 2.81 μm^2^ (mean ± S.D.) (Fig. 3F). The area of micronuclei containing mis-segregated chromosome 12 chromatids, was 12.45 ± 9.58 μm^2^ (mean ± S.D.). Micronuclei containing chromosome 12 were not significantly different in size from lamin B1-positive micronuclei and were significantly larger (p<0.0001) than lamin B1-negative micronuclei (Fig. 3F). Thus, micronuclei that contain chromosome 12 are expected to contain intact nuclear lamina and exhibit a reduced frequenc of rupture.

In cancer cells, micronuclei with absent, faint, or irregular lamin B1 exhibit significantly higher levels of DNA damage (*63*). To assess DNA damage in iPSC micronuclei, we used γ-H2AX immunolabeling (Fig. 3D). We quantified γ-H2AX presence in both primary nuclei and micronuclei. Approximately 50% of analyzed micronuclei were γ-H2AX-positive (Fig. 3E), consistent with our observation that ∼40% of micronuclei lacked intact lamin B1. The loss of lamin B1 compromises effective DNA damage repair (*64*, *65*). Micronuclei containing chromosome 12 were similar in size to γ-H2AX-negative micronuclei (10.41 ± 5.9 μm^2^, mean ± S.D.), and were significantly larger (p<0.0001) than γ-H2AX-positive micronuclei (3.46 ± 2.63 μm^2^, mean ± S.D.) (Fig. 3F). Thus chromosome 12 chromatids were mis-segregated into larger micronuclei containing intact lamina shown to be protected from DNA damage.

We hypothesized earlier that chromosome 12 chromatids mis-segregating into micronuclei (Fig. 2D-G) originate from the bridging of chromosome 12. Most of those bridges were localized to the peripheral region of the anaphase cells, apart from the central spindle (Fig. 2A-C). This is crucial, because central spindle microtubules impede the assembly of nuclear envelope proteins upon micronuclei formation on mis-segregating chromosomes, leading to defects in envelope formation, while chromosomes mis-segregating on the spindle periphery or at poles generate micronuclei with intact lamina (*59*). Consistently, we showed that the peripheral positioning of chromosome 12 bridges correlates with the formation of larger micronuclei that maintain intact lamina and exhibit a lower propensity for rupture (Fig. 3G).

### 2.6. Micronuclei in human iPSCs fail to replicate DNA in S phase

Mis-segregated chromosomes trapped in micronuclei can lead to the generation of aneuploid daughter cells (*66*). However, studies on DNA replication in micronuclei have yielded contradictory results. Some have reported a delay in DNA synthesis and decreased replication efficacy (*67*); others have argued that micronuclei replicate normally and in synchrony with the primary nucleus (*68*).

We examined DNA replication in micronuclei by incubating cells with the thymidine analog, 5-ethynyl-29-deoxyuridine (EdU). Following a short EdU pulse, cells were fixed and labeled, revealing that approximately 60% of asynchronous iPSCs were in S phase at any given EdU treatment time. This result is in agreement with previous studies mapping cell cycle patterns of human ESCs and iPSCs (*69372*). We found that nearly all micronuclei (∼95%) failed to incorporate EdU when their primary nuclei did. We did not observe evidence of delayed replication in micronuclei at later stages of the cell cycle (Fig. 3H-I). We concluded that DNA in iPSC micronuclei rarely undergoes replication, leading to chromosomes in micronuclei entering mitosis without being replicated.

### 2.7. Transition passage iPS cells enter mitosis with a single unpaired chromosome 12 chromatid

To investigate whether micronuclei with mis-segregated chromosome 12 chromatids persist through G2 phase and may lead to the generation of aneuploid daughter cells, we analyzed the mitotic segregation of chromosome 12 and chromosome 10 in FISH studies.

Normally, each chromosome is composed of 2 identical chromatids. In mitosis, the chromosomes first align at the metaphase plate. Each chromosome has 2 centromeres, and at metaphase, when centromeres are adjoined, they most often appear as a single dot when labeled by FISH. At anaphase, the chromatids separate and move toward opposite poles. The FISH signals for each chromosome now resolve as 2 dots. In diploid cells, 2 signals for both chromosomes 10 and 12 appear at metaphase and 4 signals at anaphase (Fig. 3J). In trisomic cells, 2 chromosome 10 signals and 3 chromosome 12 signals appear in metaphase. These resolve into 4 chromosome 10 signals and 6 chromosome 12 signals in anaphase (Fig. 3K).

However, in 1.5% of 798 analyzed metaphase cells during transition passages, we observed a chromosome 12 signal positioned away from the metaphase plate (Fig. 3L). Failure to fully align at metaphase would be expected if these signals reflected an unpaired chromosome 12 chromatid. This was never seen for chromosome 10. Similarly, in 2% of 655 analyzed anaphase cells in transition passages, chromosome 12 showed 2 signals oriented to one pole and 3 to the other (Fig. 3M). This 3:2 anaphase segregation was never detected for chromosome 10.

Because of the extraordinary nature of this apparent asymmetric arrangement in mitotic cells (5 chromosome 12 chromatids), we investigated whether chromosome 12 signals originated from single chromatids or whole chromosomes using quantitative fluorescence. This approach allowed us to address two important questions. First, do the chromosome 12 signals lying apart from the metaphase plate reflect whole chromosomes or single chromatids? Second, might the 3:2 anaphase cells be explained by trisomic anaphase cells where two chromosome 12 centromeres on one side overlapped, thus generating one anaphase signal that contained two chromatids? Our data revealed that the chromosome 12 fluorescence signals lying apart from the metaphase plate were consistent with most originating from a single chromatid. Similarly, the fluorescence signals in each of the centromeres in the diploid pole of 3:2 anaphase cells were primarily due to single chromatids rather than two superimposed centromeres (Fig. 3N). These data show that at least 1% of transition passage cells enter mitosis with an abnormal chromosome 12 configuration, two intact chromosomes, and one unpaired chromatid, originating from the previous mis-segregation into the micronuclei.

### 2.8. Why chromosome 12? Short telomeres and faster proliferation

#### 2.8.1. Chromosome 12p arms create bridges in anaphase during trisomy 12 takeover

Our next focus was to address the question: why is chromosome 12 prone to this mis-segregation? Certain cancers exhibit high frequencies of anaphase chromosome bridges and aneuploidy, which are known to arise from telomere dysfunctions (*73-75*), especially affecting chromosomes with the shortest telomeres (*76*). Telomeric defects, recognized as DNA double-strand breaks (DSBs), can cause chromosome bridges when uncapped telomeres join, which explains their high frequency in cells with telomere attrition (*51*, *77-79*). Intriguingly, the shortest telomeres in humans are located on the 17p, 20q, and 12p arms (*80*), and the same chromosome arms are most frequently involved in chromosomal aberrations observed in cultured human embryonic and pluripotent stem cell lines (*26*, *29*, *32*, *33*, *35*, *81*, *82*).

Chromosome 12 exhibited a high propensity for bridging and these bridges occurred at the ends of chromosome 12 arms (Fig. 2A-C). This led us to ask which chromosome 12 arms were involved in bridging. We measured the length of p and q arms in diploid and trisomic metaphase spreads (Fig. 4A-B). The 12p arm measured 1.16 ± 0.21 µm (mean ± S.D.) and 12q 2.68 ± 0.6 µm (mean ± S.D.). 12p arms were significantly shorter than the 12q in both diploid and trisomic cells (p<0.0001). No significant differences in 12p arms were observed between diploid and trisomic cells, but 12q arms were longer in trisomic metaphase spreads (p=0.0313) (Fig. 4B).

**Fig. 4.**
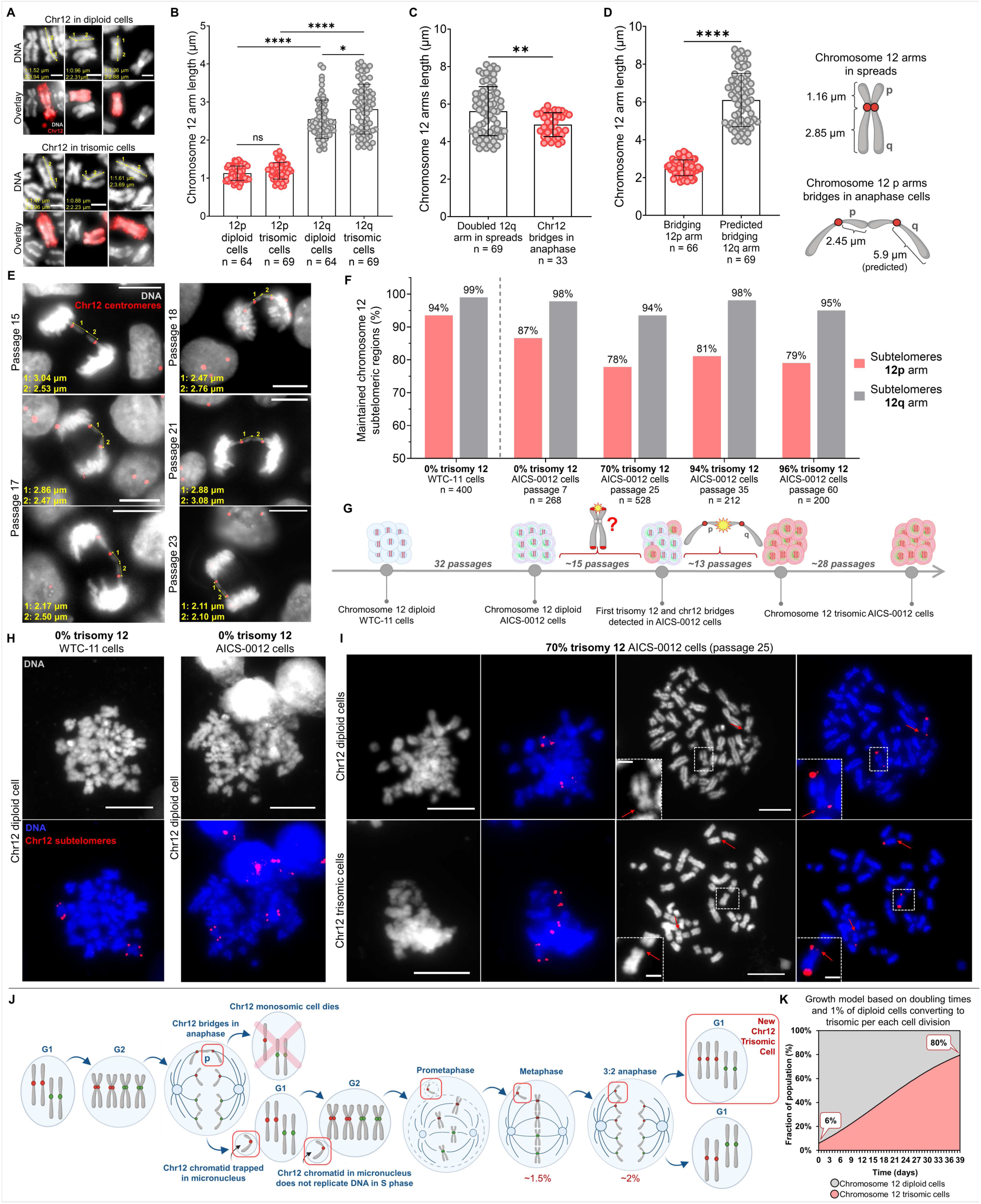
Chromosome 12p arms show a high propensity for bridging in anaphase cells and show disrupted subtelomeres, providing a pathway for the generation of trisomy 12 cells. **(A)** Chromosome 12 in diploid and trisomic metaphase spreads after labeling with whole chromosome 12 probes used to analyze p and q chromosome 12 arm length. Measurements were performed from the mid-centromeric region to the end of each arm. The overlays merge DNA (grey) and chromosome 12 (red). Scale bars = 2 µm. **(B)** 12p and 12q arms measured in diploid and trisomic metaphase spreads after labeling with the whole chromosome 12 probes. Each dot on the graph represents the length of a single arm. The n values represent the number of measured arms. Data presented as mean ± SD; p values were determined by unpaired two-tailed Student’s t-test with Welch correction and Mann-Whitney U test. P-values are as follows: * p=0.0313, **** p<0.0001. **(C)** The total length of both bridging chromosome 12 arms compared to the doubled 12q arm length in the metaphase spreads. Each dot on the graph represents the length of two arms. Data presented as mean ± SD; p value was determined by unpaired two-tailed Mann-Whitney U test, ** p=0.0097. **(D)** The length of bridging 12p arms compared to the predicted length of bridging 12q arms after applying the averaged stretch ratio of p arms. Each dot on the graph represents the length of a single arm. Data presented as mean ± SD; p value was determined by unpaired two-tailed Mann-Whitney U test, **** p<0.0001. **(E)** Examples of chromosome 12 bridges in transition passage anaphase cells used for the analysis. Measurements were performed from the mid-centromeric region to the end of each bridging arm. The overlays merge DNA (grey) and chromosome 12 centromere (red). Scale bars = 10 µm. **(F)** Percentages of retained chromosome 12 subtelomeric regions in diploid, parental WTC-11 cells, and AICS-0012 cells before, during and after trisomy 12 takeover. When trisomy 12 was gaining dominance, in passage 25 (70% trisomy 12), the loss of 12p subtelomeres was 16%, while the 12q loss was only 5.5%. These levels remained stable for at least 35 passages more as trisomy 12 reached nearly 100% of the cell population. The n values represent the number of analyzed subtelomeres on 12p and 12q arms. **(G)** Passaging history of the WTC-11 cell line during the development of the AICS-0012 cell line and further passaging of the AICS-0012 cell line through the initial appearance and takeover of trisomy 12. We hypothesize that the loss or damage of subtelomeres on 12p arms leads to the observed bridging in anaphase cells and contributes to the trisomy 12 takeover. **(H)** and **(I)** Representative images of metaphase chromosomes labeled with 12p and 12q subtelomeric probes in the diploid WTC-11 cells, and diploid and transition passages of the AICS-0012 cell line. The loss of 12p subtelomeres became maximal at transition passage 25. Red arrows indicate 12p arms lacking subtelomeric repeats and white boxes zoom on chromosomes missing 12p subtelomeres. DNA is labeled in grey and the overlays merge DNA (blue) and chromosome 12 subtelomeres (red). The white box areas were magnified 6x. Scale bars = 10 µm. Scale bars on zoomed regions = 2 µm. **(J)** Sequence of cell cycle events generating new chromosome 12 trisomic cells. When trisomy 12 takes over the culture, bridging chromosome 12 p arms creates one chromosome 12 monosomic cell that will not survive, and another daughter cell, where, in some cases, a chromosome 12 chromatid generates a micronucleus. This chromosome 12 trapped in the micronucleus fails to replicate properly in the next S phase. The resulting G2 cell has 2.5 copies (5 chromatids) of chromosome 12. This would generate a metaphase cell with one unpaired chromatid, and a 3:2 anaphase, both of which we observed in the transition passages. This 3:2 anaphase would generate a new chromosome 12 trisomic cell. **(K)** Trisomy 12 outgrowth modeled by incorporating the doubling time differences and an additional variable of 1% of diploid cells converting to trisomic at each cell division. After applying this variable, the mathematical model showed 80% of trisomy 12, closer to the observed trisomy 12 dominance.

Next, we examined the length of bridging chromosome 12 arms (Fig. 4C-E). We found that the single bridging arm was 2.45 ± 0.36 µm (mean ± S.D.). We compared the observed length of bridging arms to the doubled 12q arms in spreads. Our analysis showed that the bridging arms are 12p (p=0.0097) (Fig. 4C). We further calculated the stretch ratio of 12p arms and applied it to 12q arms. This simulation showed that if the bridging arms were 12q, their length would be approximately 5.9 µm, more than double the observed length (Fig. 4D). Thus, we confirmed that the 12p arms were forming bridges.

#### 2.8.2. During trisomy 12 takeover almost 20% of chromosome 12p arms appear with lost subtelomeric repeats

Subtelomeric sequences, located near the ends of chromosome arms and adjacent to telomeres, play a crucial role in protecting telomeres from degradation. These regions are hotspots for DNA breakage and contribute to large copy number variations (*42*, *80*, *83-86*). Long noncoding telomeric RNA (TERRA) promoters are localized at subtelomeres and are critical for maintaining telomere stability. Cells lacking CTCF binding sites within the TERRA promoter are sensitive to replicative stress and exhibit impaired DNA replication, with the formation of ultra-fine anaphase bridges during replicative stress conditions (*87-89*). This led us to hypothesize that the bridging of chromosome 12 might be due to subtelomere damage leading to telomere shortening on the 12p arms.

We labeled subtelomeres on the 12p and 12q arms in the metaphase spreads of both, the diploid, parental WTC-11 cell line and the diploid, transition, and trisomic passages of the AICS-0012 cell line (Fig. 4F-I). In chromosome 12 diploid WTC-11 cells, 94% of 12p and 99% of 12q subtelomeric regions were present. After 34 passages (development of AICS-0012 cell line from the WTC-11 parental cells), we observed a 7% decrease in 12p compared to only a 1% decrease in 12q subtelomeres. Interestingly, when comparing AICS-0012 cells with 0% trisomy 12 to those with 70% trisomy 12, we detected an additional 9% decrease in the presence of 12p subtelomeres. This decrease occurred within only 18 passages. Overall, 12p subtelomeres loss was 16%, compared to a 5% loss of 12q subtelomeres. Importantly, a substantial 9% of 12p loss occurred while cells were developing trisomy 12. Surprisingly, trisomic passages showed no further subtelomere loss beyond the decrease observed at 70% trisomy 12. From passage 25 onward, as trisomy 12 reached nearly 100% of the cell population over the following 35 passages, the level of preserved subtelomeric regions remained stable (Fig. 4F).

Our findings show that subtelomeric repeats loss or damage on the 12p arms during the critical, transition passages, probably occurring due to replication stress (*80*, *83*, *84*), can play a role in the onset and dominance of trisomy 12. This loss can disrupt the proper separation of 12p arms during anaphase, resulting in the observed bridging. The elevated propensity for subtelomere loss in the 12p arms, along with the high incidence of bridging in these arms, supports a link between subtelomeric instability and the increased rates of chromosome 12 missegregation observed during transition passages.

### 2.9. Mechanisms driving trisomy 12 takeover in human iPSCs

Based on our findings, we propose a model for how chromosome 12 trisomic cells arise in human iPSCs (Fig. 4J). From a diploid parent, bridging chromosome 12 p arms create a monosomic daughter cell that rapidly dies, and a diploid cell with an extra chromatid 12 trapped in a micronucleus. This micronucleus fails to replicate DNA during S phase and creates a G2 cell with 5 chromosome 12 chromatids. Therefore, in the next mitosis, this G2 cell generates 3:2 anaphase. The cell receiving three copies of chromosome 12 becomes a new trisomic cell.

Based only on the doubling times difference, we would have expected only 38% of trisomic cells after 13 passages (39 days in the cell culture) (Fig. 1E). However, our FISH experiments consistently showed almost 100% of trisomy 12 (Fig. 1F). Based on mathematical modeling, we hypothesized earlier that to reach almost 100% of trisomy 12 in 13 passages, 3% of diploid cells have to convert to trisomic during each division (Fig. 1G). To verify the previous model, we introduced a variable representing the percentage of diploid cells converting to trisomic per each cell division observed in our FISH mitosis studies. Our data showed that 2% of anaphase cells in transition passages exhibited 3:2 segregation, generating one diploid and one trisomic cell (Fig. 3M). Using a more conservative rate of 1% resulted in the trisomy 12 reaching a predicted 80% after 13 passages (Fig. 4K). This prediction more closely matches the observed outgrowth of trisomic cells in FISH experiments (Fig. 1F).

Overall, our data indicate that chromosome 12 trisomic cells, once present in the culture, outgrow their diploid precursors due to a faster growth rate. However, we provide evidence that this rapid takeover is driven by the simultaneous conversion of at least 1% of diploid cells to trisomic during each division. These combined mechanisms drive the rapid dominance of chromosome 12 trisomic population in human iPS cells.

## 3. Discussion

Mapping pathways by which diploid cells become aneuploid in culture is challenging because primary cultures have limited lifespans due to replicative senescence (*4*, *5*). Moreover, nearly all immortalized cell lines, including those thought to reflect <normal= cells, show some extent of chromosomal abnormality (*8-13*). Consequently, most studies of aneuploidy have relied on non-isogenic tumor-derived or transformed cell lines, making it difficult to study aneuploidy in its native context (*10*, *14*, *15*).

Human induced pluripotent stem cells (iPSCs) offer a unique opportunity to study the rise of aneuploidy from diploid cells that can proliferate indefinitely (*19*, *20*). Over time, iPSCs may acquire non-random chromosomal abnormalities, with trisomy 12 being one of the most frequent and significant changes (*25-28*, *90*). The frequent occurrence of this trisomy poses serious problems for the use of iPSCs in basic research of developmental pathways. It also endangers their use in therapeutic medicine (*27*, *34*, *35*, *91*, *92*) given that trisomy 12 is characteristic of various malignancies, including chronic lymphocytic leukemia (*90*, *93*) and germ cell tumors (*94*, *95*).

Our study aimed to address a key question: how and why does chromosome 12 trisomy arise and rapidly dominate human iPSC cultures? Previous reports have attributed this to a growth advantage conferred by trisomy 12 cells (*31*, *33*, *35-38*), but our findings indicate that growth advantage alone cannot account for the speed at which trisomic cells overtake the diploid. We provide new evidence that chromosome 12 trisomy arises through a combination of enhanced proliferation of trisomic cells and chromosome 12-specific mitotic missegregation events occurring at an unexpectedly high frequency per each cell division.

We highlighted the importance of both mechanisms, growth rate differences and the high magnitude of mis-segregation events through mathematical modeling (Fig. S6). We modeled scenarios where the doubling times of diploid and trisomic cells are identical, and high diploid-to-trisomic conversion rates (1% and 2%) are applied. These simulations show that even with a 2% conversion rate, trisomic cells do not reach 70% of the population within 13 passages, emphasizing the critical role of faster growth rates in driving the takeover (Fig. S6A). We also modeled the effect of the observed doubling time difference between diploid and trisomic cells but with much lower mis-segregation rates (0.01% and 0.1%). Under these conditions, trisomic cells fail to reach even 50% of the population within 13 passages, demonstrating that the frequency of chromosome 12 mis-segregation events must be remarkably high to account for that rapid takeover (Fig. S6B). Finally, we present comprehensive models incorporating both mechanisms: the doubling time advantage of trisomic cells and high conversion rates (1%, 2%, and 3%).

Only when both factors are combined does the model replicate the observed rapid dominance of trisomic cells in culture (Fig. S6C). Together, these findings underscore that both the difference in growth rates and the exceptionally high missegregation frequency are crucial for driving the rapid emergence and dominance of trisomy 12 in human iPSCs.

While chromosome mis-segregation errors have been well-studied in human stem and cancer cells, our use of dual-color fluorescence in situ hybridization (FISH) to track chromosome 12-specific mis-segregation reveals insights not previously captured. To the best of our knowledge, we are the first to observe the 3:2 segregation pattern of chromosome 12 in anaphase and the formation of chromosome 12 p-arm bridges. Given the unprecedented nature of the existence of a single, unpaired, and unreplicated chromosome 12 chromatid, we devised a method for quantitative fluorescence measurements of centromere FISH probes to distinguish between single chromatids and whole chromosomes. Moreover, although chromosome 12 is average in size and gene content (*40*), it forms bridges that localize to the periphery of anaphase cells, in contrast to the general tendency of primarily the largest chromosomes to segregate toward the cell’s exterior during mitosis (*96*, *97*).

Our findings underscore the important role of micronuclei in the trisomy 12 takeover. Chromosome 12 chromatids mis-segregating during mitosis frequently become trapped in micronuclei, which we confirmed by fluorescence intensity analysis as being predominantly single chromatids rather than whole mitotic chromosomes. The presence of chromosome 12 in micronuclei, particularly during transition passages, suggests that these bridging chromatids play a role in driving chromosome instability. Additionally, peripheral positioning of mis-segregating chromosome 12, the size of micronuclei containing chromosome 12 along with their lamin B1 retention, indicate that these micronuclei are more stable and less likely to rupture (*56*, *58-62*). This is significant, because micronuclei rupture is a known trigger for chromothripsis, a catastrophic genomic rearrangement, and the structural stability of these micronuclei may reduce the likelihood of further genomic damage (*53*, *55*, *57*, *98*). However, our data suggest that at least the larger micronuclei in iPSCs may be more stable than those observed in cancer cells (*53*, *57*).

A potential limitation in our study is the contribution of other aneuploidy drivers, such as the bystander effect, where neighboring cells could influence one another through secreted factors in the culture media (e.g. exosomes) (*42*). However, given the strong evidence we found for chromosome 12-specific mis-segregation, it seems unlikely that the bystander effect is a primary cause. Additionally, the method of cell passaging can affect the accumulation of chromosomal abnormalities (*81*). Enzymatic passaging is known to increase the likelihood of genetic instability, though manual passaging does not entirely eliminate the risk of variants (*29*). Therefore, while passaging may contribute to genetic changes in iPSC cultures, it does not explain the specific dominance of trisomy 12.

An intriguing aspect of our findings is the link between subtelomeric instability and chromosome 12 bridging. We observed almost 20% loss of subtelomeric regions on the 12p arms during the onset of trisomy 12, which may be driven by DNA replication stress (*80*, *83*, *84*). Telomeres and subtelomeric regions are crucial for maintaining chromosome integrity, and defects in these regions can lead to bridge formation during mitosis (*87-89*). Among the shortest telomeres in human cells are those located on the 12p arms (*80*). These may be particularly susceptible to damage, and this may explain why chromosome 12 shows such a high propensity for bridging and mis-segregation (*73-76*). Replication stress during culture, especially at the subtelomeres, could serve as a trigger for these events, leading to the onset of trisomy 12 across multiple cells simultaneously. However, telomerase in human iPS cells remains active and it maintains telomere integrity, mitigating replication stress and preventing extensive DNA damage associated with aneuploidy. This allows aneuploid cells to bypass senescence or apoptosis, ensuring their survival and proliferation despite chromosomal imbalances (*83*). Telomerase can even initiate de novo telomere synthesis at any DSBs, creating neotelomeres that heal chromosome breaks (*99*). Genomic analyses of hundreds of cancer cell lines identified telomeric insertions at various chromosomal sites, including specific regions on 12p arm (*100*). But even when telomerase is insufficient to resolve telomeric damage, alternative mechanisms such as the alternative lengthening of telomeres (ALT) pathway can be activated (*101*). This dual capacity underscores the resilience of iPSCs in managing telomeric and chromosomal instability, even under conditions of rapid aneuploidy development.

In our study, we observed that during transition passages, more than 50% of detected anaphase bridges involved chromosome 12 p arms. However, once trisomy 12 became dominant, the tendency for chromosome 12 to form bridges significantly decreased. This decline in bridging coincided with the stabilization of subtelomeric regions on 12p, as evidenced by the absence of further loss of subtelomeric repeats after cells reached full trisomy 12. At 70% trisomy 12, nearly 20% of 12p subtelomeric repeats were lost, suggesting that rapid aneuploidy progression might have temporarily overwhelmed telomeric maintenance mechanisms. However, the stabilization observed in trisomic passages suggests that telomerase and/or ALT pathways eventually restored telomere integrity. Thus, we hypothesize that the observed reduction in chromosome 12 bridges in trisomic passages reflects the successful resolution of critically short telomeres on the 12p arm. Telomerase, or possibly ALT mechanisms, likely contribute to the repair of these regions, preventing the formation of anaphase bridges and stabilizing the trisomic population. The unique challenges posed by the 12p arm, such as its short telomeres, may have initially driven the high frequency of missegregation events. However, once telomeric integrity was restored, these bridging events subsided, enabling the trisomy 12 cells to dominate the culture.

Here we show that trisomy 12 appearance and dominance in iPSC culture does not arise from rare, aberrant cells, but stems from massive mitotic mis-segregation specific to chromosome 12. The newly formed trisomic cells subsequently outgrow the diploid population. While chromosome mis-segregation, micronucleation, and telomere instability are commonly observed in various malignancies (*51-53*, *73-79*), our findings reveal their roles in inducing chromosome 12 trisomy in human stem cells. By uncovering de novo emergence of trisomic cells, bridging exclusively through 12p arms, unpaired chromosome 12 chromatids in mitosis, and subtelomeric instability during transition passages, we provide new insights into trisomy 12 dominance. Our discovery represents a novel paradigm for whole chromosome instability, with significant implications for research and regenerative therapies involving stem cells, as well as for understanding the origins of aneuploidy.

## 4. Materials and Methods

### 4.1. Human induced pluripotent stem cell lines

Human iPS cell lines used in this study, identifiers AICS-0012 cl. 105 (TUBA1B-mEGFP) and AICS-0013 cl. 210 (LMNB1-mEGFP) were derived from the WTC-11 cell line at the Allen Cell Institute. WTC-11 cell line has been derived from healthy, male donor fibroblast cells using episomal vectors. The identity of the unedited WTC-11 line was confirmed with short tandem repeat profiling. Whole genome sequencing and population-level RNA-seq data for the WTC-11 iPSC line are publicly available from the Allen Cell Institute/UCSC Genome Browser (*102*). The genetically edited iPSC lines were obtained from the Allen Cell Collection. Each gene-edited cell line harbors a fluorescently tagged protein localizing to a cellular structure. The CRISPR/Cas9-mediated genome editing methodology used to generate these cell lines was described in (*103*). The donor plasmids used to generate AICS-0012 cell line is AICSDP-4 TUBA1B-mEGFP (Cat. #87421) and to generate AICS-0013 cell line is AICSDP-10: LMNB1-mEGFP (Cat. #87422) and can be obtained through Addgene (https://www.addgene.org/The_Allen_Institute_for_Cell_Science/). Specifically, to obtain the AICS-0012 cell line, using CRISPR/Cas9, WTC-11 iPSCs at passage 33 were endogenously tagged with mEGFP in a single allele of the *TUBA1B* gene and then cloned (clone 105) at the Allen Cell Institute. AICS-0012 cells were obtained from the Coriell Institute at passage 32 from the Allen Cell Collection and then cultured in this study between passages 1 to 61. To obtain the AICS-0013 cell line, using CRISPR/Cas9, WTC-11 iPSCs at passage 33 were endogenously tagged with mEGFP in a single allele of the *LMNB1* gene and then cloned (clone 210) at the Allen Cell Institute. AICS-0013 cells were obtained from the Coriell Institute at passage 30 from the Allen Cell Collection and then cultured in this study between passages 1 to 16. Edited AICS-0012 and AICS-0013 cell lines were karyotypically normal based on at least 20 metaphase cells analyzed by G-banding, did not possess any mutations at off-target sites, and had no acquired mutations 8 passages beyond the original line WTC-11 that affect genes in Cosmic Cancer Gene Census. WTC-11 is the only cell line used by the Allen Institute. Cell lines are described at https://www.allencell.org and available through Coriell at https://www.coriell.org/1/AllenCellCollection.

### 4.2. Human induced pluripotent stem cell culture

Human iPS cell lines were cultured in 25 cm^2^ and 12.5 cm^2^ flasks during standard passaging, in 96-well plates for proliferation assays, 6-cm cell culture dishes for chromosome spreads and in 6-well plates on square cover glass coverslips for FISH and immunofluorescence assays. All grow surfaces were coated in Growth Factor Reduced (GFR) Matrigel (Cat. #354230, Corning) diluted in phenol red-free DMEM/F12 (Cat. #21041025, Gibco, Thermo Fisher Scientific) at the ∼1:30 ratio (Matrigel final protein concentration = 0.337mg/mL). Matrigel plates and flasks were stored at 4°C and used within 1 week from coating.

Cells were maintained with mTeSR Plus media (Cat. #1000276 and #1000274, STEMCELL Technologies) supplemented with 1% penicillin-streptomycin (Cat. #SV30010, Cytiva). Cells were passaged every 3 days (∼72 hours) as single cells. At each passage, iPSCs were split between a 1:4 to 1:8 ratio depending on density, the following experiment design, or when colonies began to merge. Cells were transferred to a new flask following detachment via Accutase (Cat. #A1110501, Gibco, Thermo Fisher Scientific) and blocking via 1X Dulbecco’s Phosphate-Buffered Saline (DPBS) (Cat. #14190144, Gibco, Thermo Fisher Scientific) according to manufacturer’s instructions. Culturing media was supplemented with 10 μM Y-27632 Rho-associated protein kinase (ROCK) inhibitor (Cat. #HY-10583, MedChem Express) for approximately 24 hours following passaging, then changed to media without ROCK inhibitor. Cells were fed daily with fresh room temperature (RT) complete mTeSR Plus media and maintained at 37°C under a humidified atmosphere of 5% CO_2_ in a water-jacketed incubator. Cells were passaged and maintained continuously for various passaging times but a maximum of ∼180 days (61 passages). Human iPSC colonies maintained normal morphology and had an average death rate of 2%. Trypan-blue stain (Cat. #T10282, Invitrogen, Thermo Fisher Scientific) was used for cell counts and to assess the percentage of dead cells at each passage using the Countess Automated Cell Counter. A detailed protocol is available at the Allen Cell Explorer (https://www.allencell.org/cell-catalog.html, SOP: WTC culture v1.7., Allen Institute for Cell Science, 2020).

### 4.3. Fluorescent in situ Hybridization (FISH)

#### 4.3.1. Preparation of metaphase chromosomes from human iPSCs

Human iPSCs were grown in 6-cm cell culture dish until reached 80-90% confluence and were treated with 0.1 ug/mL Colcemid (Cat. #15212012, KaryoMAX Colcemid, Gibco, Thermo Fisher Scientific) and 0.5 uM Wee1 inhibitor (PD0166285, Cat. #HY-13925, MedChemExpress) for 2.5 hours. The medium was transferred to the 15 mL conical; cells were washed with 5-10 mL sterile 1X PBS (without Ca and Mg) and added to the conical. Cells were treated with Accutase, transferred to the same conical and centrifuged for 5 min at 1000 rpm. Leaving ∼0.5 mL of medium, cell pellet was gently resuspended and treated with ∼5 mL of prewarmed (37°C) 0.075M KCl swelling buffer (volume of hypotonic solution was dependent upon the size of the cell pellet), drop-by-drop, while agitating gently. Cells were incubated in a 37°C water bath for 25 min. 4-5 drops of freshly prepared Carnoy’s fixative (methanol: glacial acetic acid; 3:1 per volume basis (v/v)) was added to stop the reaction. Cells were centrifuged for 5 min at 1200 rpm and 5 mL of freshly prepared fixative was added gradually, drop-by-drop while gently agitating the tube and mixed well by flicking the tube so no clumps of cells remain.

Centrifugation and resuspension in fresh fixative were repeated twice. Microscope slides and coverslips were washed in soapy water, rinsed 3 times with Milli-Q water before use, kept in Milli-Q water at 4°C for 1 hour, and dried well right before use. Slides and coverslips were humidified at 42°C in the oven on wet paper towels for 2-3 min right before spreads preparation. Metaphase chromosomes were prepared by placing 50-150 uL (depending upon the size of the initial cell pellet) of preparation dropped onto clean glass from a height of 60 cm. Coverslips were dried overnight on a parafilm in a humidified chamber (150 mm dish on top of wet filter paper). The next day, metaphase preparations were checked for the quality of chromosome spreads using 42, 6-diamidino-2-phenylindole (DAPI) staining (1000 ng/mL). Imaging was performed with a Zeiss Axioplan II microscope platform using a 100× oil objective, Hamamatsu Orca II camera, and analyzed with the FIJI ImageJ software. Before performing FISH on metaphase preparations, coverslips were incubated at 37°C overnight.

#### 4.3.2. Dual-color FISH labeling of chromosome 10 and chromosome 12 centromeres

6-well plates with 18 mm or 22 mm square glass coverslips were coated with 1.5 mL of Matrigel and used within one week from the coating. FISH experiments for trisomy 12 takeover calculations were performed on coverslips made every 3 days alongside regular cell passaging using cells coming from the corresponding flasks. Human iPSCs were plated as a single cell suspension to assure monolayer cell growth and the confluence matching the corresponding flasks after 72 hours of growth.

Oligonucleotide DNA FISH probes designed to label alpha satellite repeats on centromeres of chromosome 10 (locus D10Z1, region 10p11.1 - q11.1) in green (Cat. #LPE 010G) and chromosome 12 (locus D12Z3, region 12p11.1-q11.1) in red (Cat. #LPE 012R), as well as hybridization buffer were purchased from the Cytocell (OGT Sysmex Group Company, Cambridge, UK). Probes were delivered as 15 uL concentrated solutions. FISH protocol was adjusted based on the official Cytocell protocol available online (https://www.ogt.com/media/za0brc0w/lpe-r-g-v012-00.pdf).

##### Day 1

Cells were grown as colonies and before fixation were rinsed with 1X DPBS. Coverslips were fixed for 15 min in Carnoy’s solution on an orbital shaker, fixative was aspirated, and coverslips were air-dried. Coverslips were rehydrated in 2X concentrated saline sodium citrate (SSC) for 2 min, dehydrated in a series of ethanol dilutions (70%, 80%, and 100%, 2 min each), and left to air-dry. Dual-color probe mix was prepared by adding 0.80 uL of each concentrated probe to 5.90 uL of hybridization buffer (7.5 uL total per 18mm coverslip). Microscope slides, coverslips, and probes solution were pre-heated on a 37°C hotplate for 5 min. Cells were then sealed with rubber glue cement with premixed probes solution in hybridization buffer, denatured on a 75°C hotplate for 2 min, and placed in a humidified lightproof container at 37°C for ∼20 h.

##### Day 2

Coverslips were washed in 0.4X SSC (pH 7.0) by immersion for 2 min at 0.4X SSC wash (pH=7.0) at 72°C and 30 s in 2X SSC/0.05% Tween 20 at RT. Coverslips were air-dried, stained with DAPI (200 ng/mL) for 5-10 min, mounted using Vectashield mounting media (Vector Laboratories, Cat. #H-1000-10 or #H-1700) and sealed with clear nail polish. Slides were stored in the dark at −20°C until image capture.

Images were taken using a Zeiss Axioplan II microscope with either 63× or 100× Zeiss oil objectives, a Hamamatsu ORCA II camera (Hamamatsu Photonics) and processed using FIJI ImageJ software. Each coverslip was initially imaged in the blue channel (DAPI). Dozens of positions were chosen at random to ensure hundreds of nuclei for trisomy 12 ratio calculations. Imaging assured the height of the z-stack capturing the entire nuclei of interphase and mitotic cells. The imaged loci were D10Z1 (chromosome 10 centromeres) labeled with fluorescein isothiocyanate and D12Z3 (chromosome 12 centromeres) labeled with Texas Red. Z stacks were taken every 200-400 nm. Before the probes were used on fixed, regularly grown cell colonies, the localization of FISH probes to the centromeres was confirmed on metaphase preparations.

##### Trisomy 12 calculations

Trisomy 12 ratio in iPSCs was calculated based on FISH performed on fixed cell colonies from multiple passages. Chromosome 10 and chromosome 12 signals were counted in hundreds of nuclei per passage. Centromeric signals for each chromosome to define trisomy 12 percentage were identified from maximal z projections of the green (chromosome 10) and red (chromosome 12) channels, followed by the ImageJ’s find maxima selection. Choosing points as an output type and the level of prominence, points9 selection represents point counts of chromosome 10 and chromosome 12 in each microscope field. ∼100% cell population diploid for chromosome 12 resulted in an average chromosome 12 to chromosome 10 ratio of ∼1.0. ∼100% trisomic cell population for chromosome 12 resulted in a ratio of chromosome 12 to chromosome 10 of ∼1.5. Counts for each pair of red and green signals within a microscope field were calculated on an unbiased spot-basis manner before each image was merged with its corresponding DNA stain. There is no mention of whole chromosomal gains or losses of chromosome 10 used as a control, in human iPS or ES cells (*29*, *31*, *41-46*). Consequently, the number of cells in each microscope field was determined by dividing the total count of chromosome 10 centromeres by two. In all FISH experiments, chromosome 10 labeling consistently indicated a diploid cell population. We acknowledge that the ratio calculations might be subject to slight errors due to imperfect labeling of the centromeric probes. Variations in the intensities of chromosome 10 or chromosome 12 centromeres labeling could potentially shift the ratio. To minimize these possible errors, we analyzed hundreds of cells from different microscope fields per passage.

##### Fluorescence intensity

The integrated fluorescence intensity of chromosome 12 centromeric regions in micronuclei, mitotic and interphase cells was measured from z-stack images and calculated relative to the background fluorescence. Images were first converted to maximum projections and then to sums using Metamorph software. The F.I. of two chromosome 12 centromeres adjoined in metaphase cells was normalized to a value of 2. Chromosome 12 F.I. in micronuclei, other mitotic and interphase cells was normalized to the F.I. of chromosome 12 centromeres in metaphase cells from the same sample. F.I. measurements and comparison were assessed only for images with the same z-stack distance and exposure time imaged during the same imaging session to maintain uniformity of fluorescence intensity measurements.

#### 4.3.3. Whole chromosome 12 labeling

FISH labeling of the whole chromosome 12 was performed on metaphase spreads and on cells grown in colonies prepared as described in 4.3.1. and 4.3.2. Probes designed to label the whole chromosome 12 with Rhodamine were purchased from ASI (Cat. # FPRPR0167, Applied Spectral Imaging Inc., CA, US) and delivered as a ready-to-use complete paintbox set. FISH protocol was adjusted based on the delivered ASI protocol.

##### Day 1

Coverslips were rehydrated in 2X SSC at RT for 2 min and then dehydrated in ethanol series: 70%, 80% and 100%, 2 min and air-dried. Then they were placed in the denaturation solution (70% formamide/2X SSC, pH = 7) at 72°C for 1.5 min. Coverslips were immediately placed in ice-cold 70%, RT 80%, and RT 100% ethanol, 2 min each and air-dried. 7.5 uL of the probe mixture per 18 mm coverslip was denatured by incubation at 80°C in a water bath for 7 minutes and then at 37°C for 10 min. Chromosome preparations were sealed with rubber glue cement with 7.5 uL of probe solution, denatured on a 74°C hotplate for 4 min and placed in a humidified lightproof container at 37°C for ∼20 h.

##### Day 2

Excess probe was removed by washing for 4 min in 0.4X SSC stringent wash (pH=7.0) at 74°C and for 2 min s in 4X SSC/0.1% Tween 20 at RT. Coverslips were stained with DAPI (1000 ng/mL) for 5-10 min, and mounted using Vectashield mounting media (Vector Laboratories, Cat. #H-1000-10 or #H-1700) and then sealed with clear nail polish. Slides were stored in the dark at −20°C until image capture.

Imaging on chromosome 12 was performed using a Zeiss Axioplan II microscope with 100× Zeiss oil objectives, a Hamamatsu ORCA II camera (Hamamatsu Photonics) and processed using FIJI ImageJ software. Each coverslip was initially imaged in the blue channel (DAPI). Z stacks were taken every 200-400 nm. When used on cell colonies, in mitotic cells, probes facilitated clear distinction between two or three chromosome 12 copies, in interphase cells, the probes exhibited the expected scattered distribution. <u>Chromosome 12 arm length</u>: Chromosome 12 in diploid and trisomic metaphase spreads after labeling with whole chromosome 12 probes were used for p and q arms9 length analysis. Measurements were performed using FIJI ImageJ software using the straight-line tool from the mid-centromeric region to the end of each arm on single-plane images of the spread chromosomes. For bridging chromosome 12 arms, measurements were also taken from the mid-centromeric region to the end of each arm, after labeling with chromosome 12 centromeric probes. These length measurements were conducted on single-plane DNA images merged with single-plane chromosome 12 centromere images to ensure clear visibility of the ends of the bridging arms and the localization of centromeres.

#### 4.3.4. Labeling subtelomeres on p and q arms of chromosomes 12

Oligonucleotide DNA FISH probes designed to label in red (Cy3 spectra) subtelomeres on p (Cat. #LPT12PR) and q (Cat. #LPT12QR) arms of chromosome 12, as well as hybridization buffer, were purchased from the Cytocell (OGT Sysmex Group Company, Cambridge, UK). Cytocell’s subtelomere probes label the most distal region of chromosome 12-specific DNA. Beyond this unique sequence is only the region of telomere-associated repeat followed by the cap of tandemly repeated (TTAGGG)_n_ sequence. FISH was performed on metaphase chromosome spreads prepared as described in 4.3.1. The FISH protocol was performed as the official Cytocell’s protocol available online: https://www.ogt.com/media/zc5bqjcb/lpt-r-g-v012-00.pdf. Imaging was performed as described in 4.3.3. <u>Assessment of subtelomeric repeats loss:</u> To evaluate the presence or loss of subtelomeric regions, the F.I. of labeled 12p and 12q subtelomeric regions was analyzed. The integrated F.I. of chromosome 12 p and q subtelomeres in metaphase spreads was measured from z-stack images and calculated relative to the background fluorescence. These images were first converted to maximum projections and then to sums using Metamorph software. Subtelomeric labeling was considered present if the F.I. of the probes was g 20% relative to the average F.I. of other labeled chromosome 12 arms within the same metaphase preparation. F.I. for each subtelomeric region was normalized to 1. If the normalized F.I. of a labeled arm was <0.2, the subtelomeric region was classified as missing. F.I. normalization and comparisons were conducted under identical imaging conditions, with all images acquired using the same z-stack interval and exposure settings within each imaging session to ensure measurement consistency.

### 4.4. EdU pulse labeling

Human iPSCs from early transition passages 13 and 14 (20-30% of trisomy 12) were seeded on 22 mm coverslips in 6-well plates. Cells were initially cultured for 24 hours in media supplemented with ROCK inhibitor. The following day, the media was replaced with regular mTeSR Plus, and cells were allowed to grow until day 3. On day 3, cells were pulsed with 20 μM EdU (5-ethynyl-29-deoxyuridine, Life Technologies, Cat. #A10044) a nucleoside analog of thymidine, for varying time intervals (10, 15, 20, and 30 min). After EdU incorporation, the media was removed, and cells were washed with 1X PBS. Cells were fixed in 1.5% paraformaldehyde (PFA)/1X PHEM buffer (60 mM PIPES, 25 mM HEPES, 10 mM EGTA, and 4 mM MgCl_2_) for 15 minutes, rinsed with 1X PHEM buffer, and permeabilized with 1X PHEM buffer containing 0.5% Triton X-100 for 8 minutes. Subsequently, cells were washed with 1X PBS. EdU was detected using click chemistry. Coverslips were incubated with the reaction mixture containing 2 mM CuSO_4_, 50 mM ascorbic acid, and 2 mM Alexa Fluor 555 Azide (Life Technologies, Cat. #A20012) for 30 minutes at RT in the dark. Incubation step was repeated twice. Following the staining, coverslips were washed twice with 1X PBS for 10 min each. Coverslips were stained with DAPI (500 ng/mL) for 10 min, mounted using Vectashield mounting medium (Vector Laboratories, Cat. #H-1000-10 or #H-1700), and sealed with clear nail polish. Prepared slides were stored in the dark at −20°C until imaging. Images were acquired using a Zeiss Axioplan II microscope with a 100× Zeiss oil immersion objective and a Hamamatsu ORCA II camera (Hamamatsu Photonics). Z-stacks were taken every 200-400 nm. Image processing and analysis were performed using FIJI ImageJ software. EdU foci presence in primary nuclei and their micronuclei were assessed. All mitotic cells were EdU negative, and at each time point of Edu treatment, approximately 60% of labeled cells were in the S phase, aligning with literature data regarding cell cycle timing in human iPSCs (*69-72*).

### 4.5. GFP visualization of lamin B1 in fixed LMNB1-GFP cells

Human iPS cell line AICS-0013, which has an mEGFP tag on the N-terminus of the lamin B1 gene, was cultured from passage 1 to 16. Cells from passages 12-14 (corresponding to the early transition passages in the AICS-0012 cell line) were used to assess lamin B1 expression in micronuclei of human iPSCs. Cells were grown on 22 mm coverslips until they reached approximately 80% confluence. Fixation was performed using 1.5% paraformaldehyde (PFA) in 1X PHEM buffer, followed by extraction using 1X PHEM, 0.5% Triton X-100, and 1:1000 protease inhibitors. After fixation for 15 minutes, cells were incubated in the extraction buffer for 8 minutes. Post-extraction, coverslips were rinsed in 1X PHEM, stained with DAPI (500 ng/mL) for 10 minutes, mounted using Vectashield mounting medium (Vector Laboratories, Cat. #H-1000-10 or #H-1700) and sealed with clear nail polish. Prepared slides were stored at −20°C in the dark until imaging. Imaging was performed using a Zeiss Axioplan II microscope with a 63× and 100× Zeiss oil immersion objective and a Hamamatsu ORCA II camera (Hamamatsu Photonics). Z-stacks were taken every 200-400 nm. Image processing and analysis were performed using FIJI ImageJ software. 3D projection analysis to assess lamin B1 integrity in micronuclear membranes was performed using MetaMorph software.

### 4.6. γ-H2AX immunolabeling

Human iPSCs from late transition passages were grown on 18 mm coverslips. Cells were pre-permeabilized in ice-cold 1X PHEM/0.5% Triton X-100/1:1000 PI for 5 min and washed twice with ice-cold 1X PBS for 5 min each. The cells were then fixed in freshly dissolved 1.5% paraformaldehyde (PFA) in 1X PHEM for 15 min at RT. Following fixation, the cells were permeabilized in 0.2% Triton X-100 in ice-cold 1X PBS for 10 min. Blocking was performed using 20% BNGS (boiled normal goat serum) in 1X MBST (MOPS-buffered saline with 0.05% Tween 20) for 30 min. Coverslips were incubated with primary antibody (monoclonal mouse anti-phospho-histone H2A.X (Ser139) (Sigma Millipore, Cat. #05-363)) diluted 1:1000 in 5% BNGS/1X MBST, for 2.5 hours. After primary antibody incubation, coverslips were washed twice with 1X MBST for 5 min each and then incubated with secondary Cy3 goat anti-mouse antibody (Jackson ImmunoResearch, Cat. #115-165-146) diluted 1:800 in 5% BNGS/1X MBST, for 1 hour in a dark, humidified chamber. Following secondary antibody incubation, coverslips were washed twice with 1X MBST for 5 min each. The DNA was labeled with DAPI (500 ng/mL) for 10 min, mounted using Vectashield mounting medium (Vector Laboratories, Cat. #H-1000-10 or #H-1700), and sealed with clear nail polish. A negative control was processed identically to the test samples, omitting primary antibodies. The test samples were analyzed and quantified relative to the negative control, which was used to standardize image scaling. Prepared slides were stored at −20°C in the dark until imaging. Imaging was performed using a Zeiss Axioplan II microscope with a 63× and 100× Zeiss oil immersion objective and a Hamamatsu ORCA II camera (Hamamatsu Photonics). Z-stacks were taken every 200-400 nm. Image processing and analysis were performed using FIJI ImageJ software.

### 4.7. Micronuclei area measurements

To determine the sizes of micronuclei, z-stack images were converted to the maximum projected areas. Micronuclei displayed an elliptical shape, and their areas were calculated using the formula for an ellipse: area [μm^2^] = π × (length/2) × (width/2). Micronuclei sizes were evaluated in human iPSCs across three experimental protocols, as detailed in sections 4.3.2, 4.5, and 4.6. Thus, the primary nuclei sizes were measured first, with 100 nuclei analyzed per protocol, across at least three independent samples for each condition. Nuclei sizes obtained following the Carnoy’s fixation (methanol: glacial acetic acid; 3:1 (v/v)) in the dual-color FISH protocol (section 4.3.2) were found to be larger than those observed following γ-H2AX immunolabeling protocol (section 4.6) and LMNB1-GFP fixation (section 4.5). To account for this variation, nuclei sizes measured from the dual-color FISH protocol were normalized to a value of 1. Correction factors were subsequently applied to the nuclei sizes obtained from γ-H2AX immunolabeling and GFP-LMNB1 visualization to align with the FISH-derived measurements. These correction factors were then applied to the micronuclei area measurements for both γ-H2AX-labeled and LMNB1-GFP expressing cells.

### 4.8. Microwell growth assay

The microwell growth assay was used to determine the doubling times of chromosome 12 diploid and trisomic cell populations. SYBR Gold nucleic acid stain (10 000X in DMSO, Cat. #S11494, Invitrogen) was used to measure fluorescence intensity based on the DNA content.

#### Cell seeding and preparation

Cells were cultured in 96-well black-walled plates with a clear, flat bottom (Cat. #655090, Greiner Bio-One). The wells were pre-treated with Matrigel (100 µL/well) the day before seeding. Matrigel was then removed, and cells were seeded in a 100 µL cell suspension per well with media containing ROCK inhibitor for the first 24 hours. Negative control wells pre-treated with Matrigel and containing only media (with and without ROCK inhibitor) were included on each plate in eight technical replicates to account for background fluorescence.

#### Standard growth curves

To establish a correlation between cell counts and fluorescence intensity, cells from diploid and trisomic populations were incubated for 3-4 hours to allow attachment. The media was removed, and cells were fixed and stained with 1% PFA, 1X PHEM, 1:1000 SYBR Gold, and 0.5% Triton X-100 for 30 min at RT. Fluorescence intensity was measured using a microplate reader (Tecan Phenix GENios, Switzerland) with consistent gain settings to ensure comparability across experiments. Measurements were taken from the bottom of the wells, using excitation and emission wavelengths of 485 nm and 535 nm, respectively. Diploid and trisomic cell populations were examined in three biological replicates, with varying cell counts (800, 1600, 3200, 6400, 12800, 26500, 51200, and 102400 cells) seeded in eight technical replicates per count. Fluorescence intensities were then used to generate linear equations from power trendlines for each population.

#### Doubling times calculation

The linear equations for both cell populations were used to calculate the end-point cell counts during the growth assays. Two types of microwell growth assays were conducted on diploid passages and trisomic passages. Time-course growth assay: Five different cell counts (ranging between 400 to 2000 cells) were seeded in four technical replicates per count, and end-point fluorescence was measured after 12, 24, 49, 74, and 96 hours of growth. Single time-point growth assay: Varying cell counts (ranging between 500 to 4000 cells) were seeded per microwell in four technical replicates, and end-point fluorescence was measured after 70 hours. End-point cell counts were calculated using the linear equations from standard growth curves. Doubling times were determined using the known seeded cell counts, calculated end-point cell counts, and the growth duration. All doubling time data were analyzed using the ROUT method (Q = 5%) to identify and exclude outliers. Final doubling times were calculated from the cleaned dataset after removing identified outliers.

### 4.9. Predictive mathematical modeling

A mathematical model created using Microsoft Excel predicted fractions of cells within a culture containing a mixed population of diploid and trisomic cells, exhibiting unique characteristics. The model allows for manipulation of key parameters, including cell doubling times, death rates, and the rate of conversion from diploid to trisomic cells. By adjusting these parameters, the model can predict the theoretical fractions of diploid and trisomic cells within the culture at any given time, depending on the input characteristics. In the model incorporating difference in doubling times, the division rate of diploid cells is normalized to 1, while the division rate of trisomic cells is 1.09, reflecting their faster growth calculated as the ratio between diploid to trisomic cell doubling times. Additionally, the initial percentage of diploid cells in the culture is an adjustable input. The model includes a variable for the conversion rate of diploid cells to trisomic cells. In the initial, theoretical scenario where trisomic cells overtake diploid cells purely by growth rates difference, this conversion rate was set to 0%. However, in a revised model that accounts for the spontaneous conversion of diploid cells to trisomic cells, this variable was set to 1%, meaning that with every cell division, 1% of diploid cells convert to trisomic cells due to chromosome 12-specific mis-segregation error. Microsoft Excel was used to create stacked area graphs to visually represent the predicted growth dynamics of both cell populations over time, illustrating how varying the input parameters affects the outcome of the culture’s cellular composition.

### 4.10. Statistical Analysis

Statistical analysis and outliers9 identification were performed using GraphPad Prism software version 10.2.3. Star values represent the level of statistical significance, with p-values as follows: * = p < 0.05, ** = p < 0.01, *** = p < 0.001, and **** = p < 0.0001.

## Acknowledgments

We thank John Daum for help with the interpretation of results and Dr. Courtney Sansam for assistance with the EdU labeling protocol. We also thank Dr. Christopher Sansam, Kevin Boyd, MSc, the OMRF Center for Biomedical Data Sciences, and Dr. Jonathan Wren for their support in aiding in the overall execution of the project. Additionally, we thank Drs. Rafal Donczew, Elizabeth Finn, Jacob Kirkland and Christopher Sansam for careful reading of the manuscript and insightful suggestions. We additionally thank Paul Gorbsky for his assistance with the mathematical modeling. The graphical abstract and diagrammatic images were created in https://BioRender.com (accessed September 2024).

## Supplementary Materials

Figs. S1 to S3.

## Author Contributions

Conceptualization: MN, GJG; Methodology: MN, MCL; Investigation: MN, MCL; Visualization: MN; Funding acquisition: GJG; Project administration: GJG; Supervision: GJG, MN; Writing 3 original draft: MN; Writing 3 review & editing: MCL, GJG.

## Data and materials availability

All data are available in the main text or the supplementary materials.

---

**Fig. S1.**
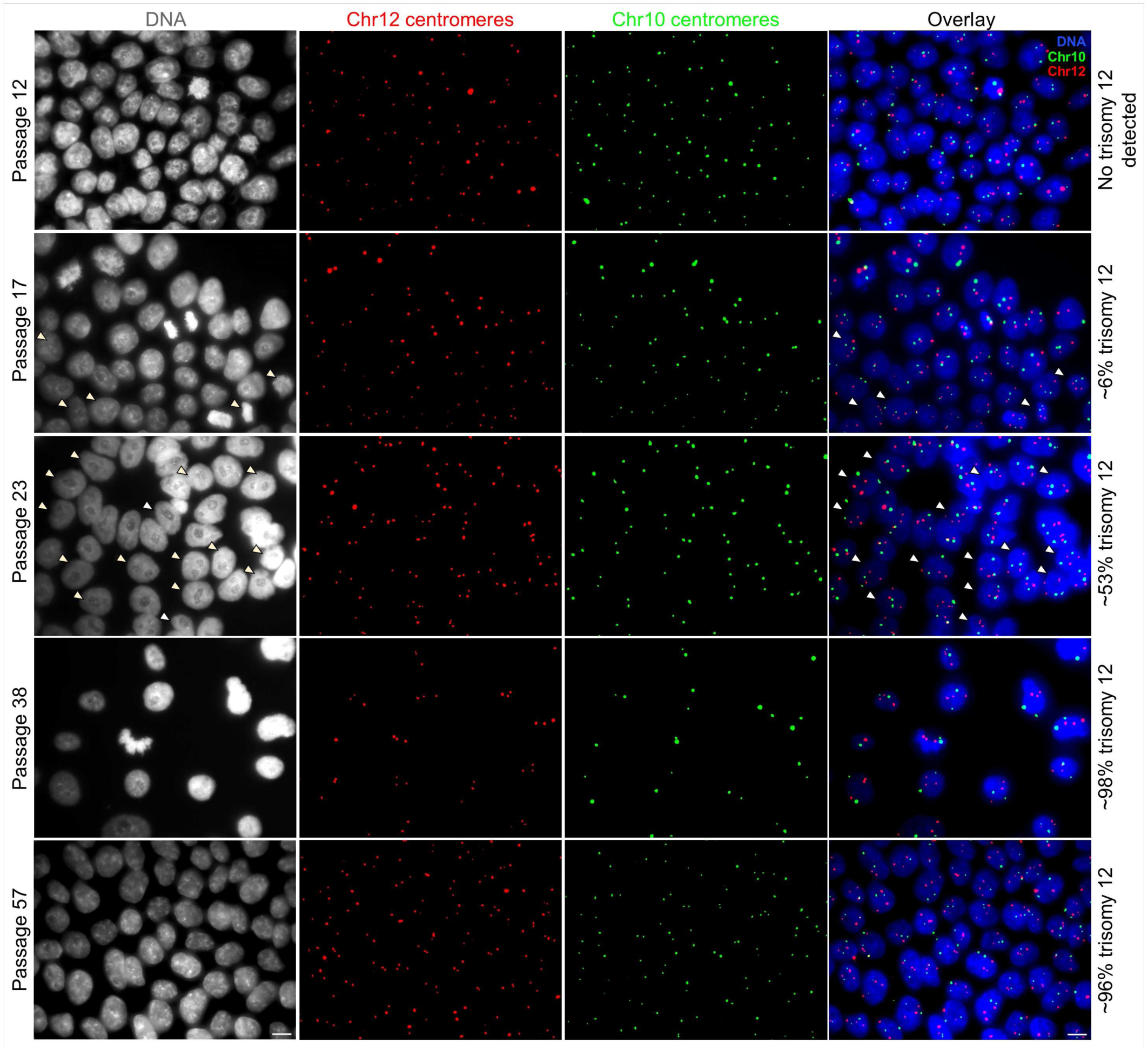
Chromosome 12 trisomic cells appear spontaneously and rise to nearly 100% of the culture during 12 passages. Representative microscope fields of iPS cells from passages 12, 17, 23, 38, and 57. In passage 12, trisomy 12 is still undetected, but in transition passages 17, and 23 we observe respectively ∼6% and ∼53% of trisomy 12. Trisomy 12 persists in the cell culture for at least 28 more post-transition passages since it gained almost 100% dominance at passage 29. Arrows on the overlays indicate chromosome 12 trisomic cells in transition passages. DNA labeling is shown in grey. The overlays merge DNA (blue), chromosome 12 centromeres (red dots), and chromosome 10 centromeres (green dots). Images for ratio calculations were taken using a Zeiss Axioplan II microscope with 63× and 100× objectives. Passage numbering reflects the number of passages performed during this study starting from passage 2 of human iPS cell line AICS-0012. Scale bar = 10 µm.

**Fig. S2.**
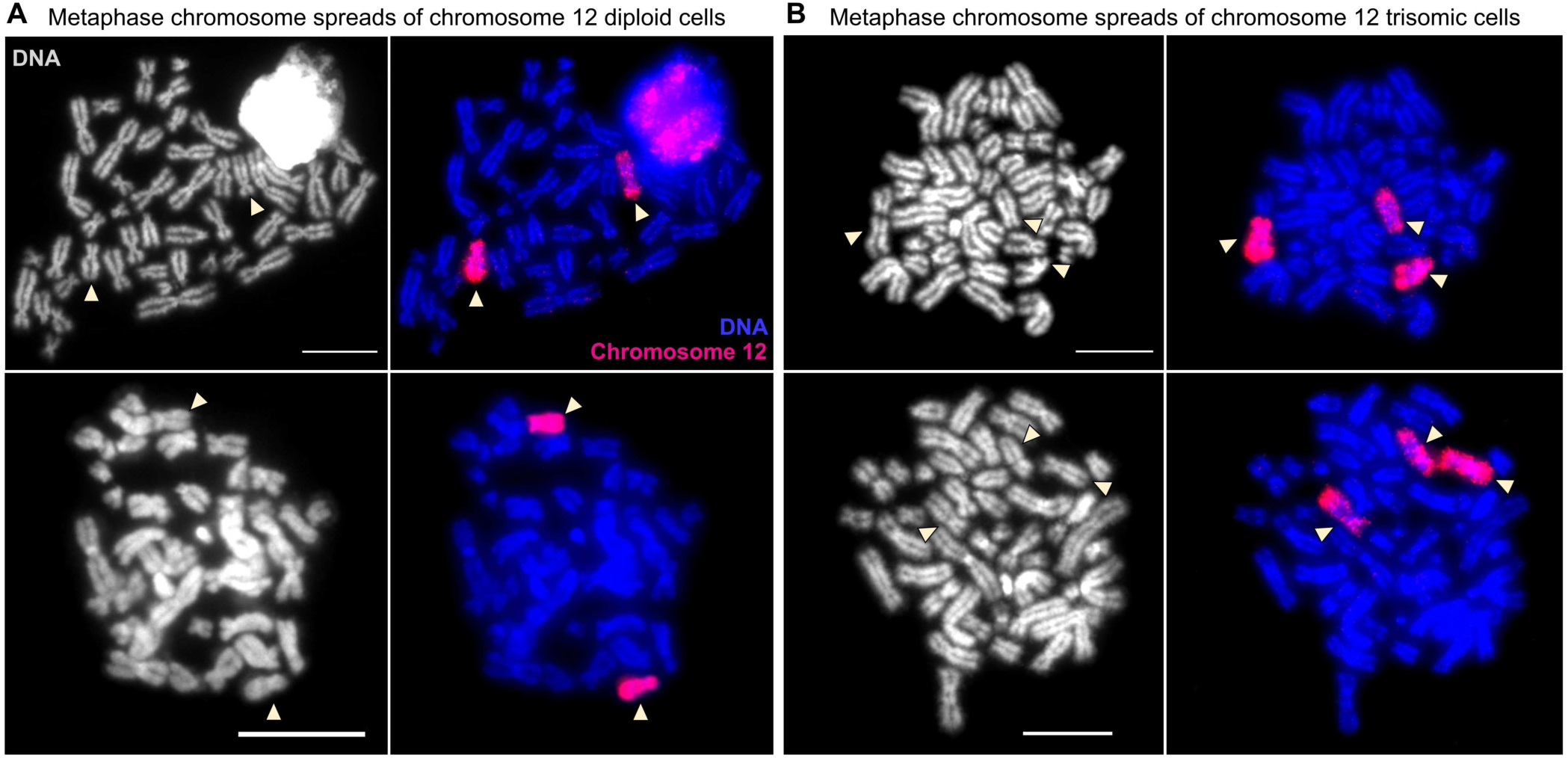
Whole chromosome 12 probes detect three copies of the entire chromosome 12 in a trisomic cell population. **(A)** Diploid metaphase spreads show two copies of chromosome 12, while **(B)** trisomic always show the entire three copies of chromosome 12. Arrows indicate chromosome 12. DNA labeling is shown in grey and the overlays merge DNA (blue) and chromosome 12 (red). Scale bars = 10 µm.

**Fig. S3.**
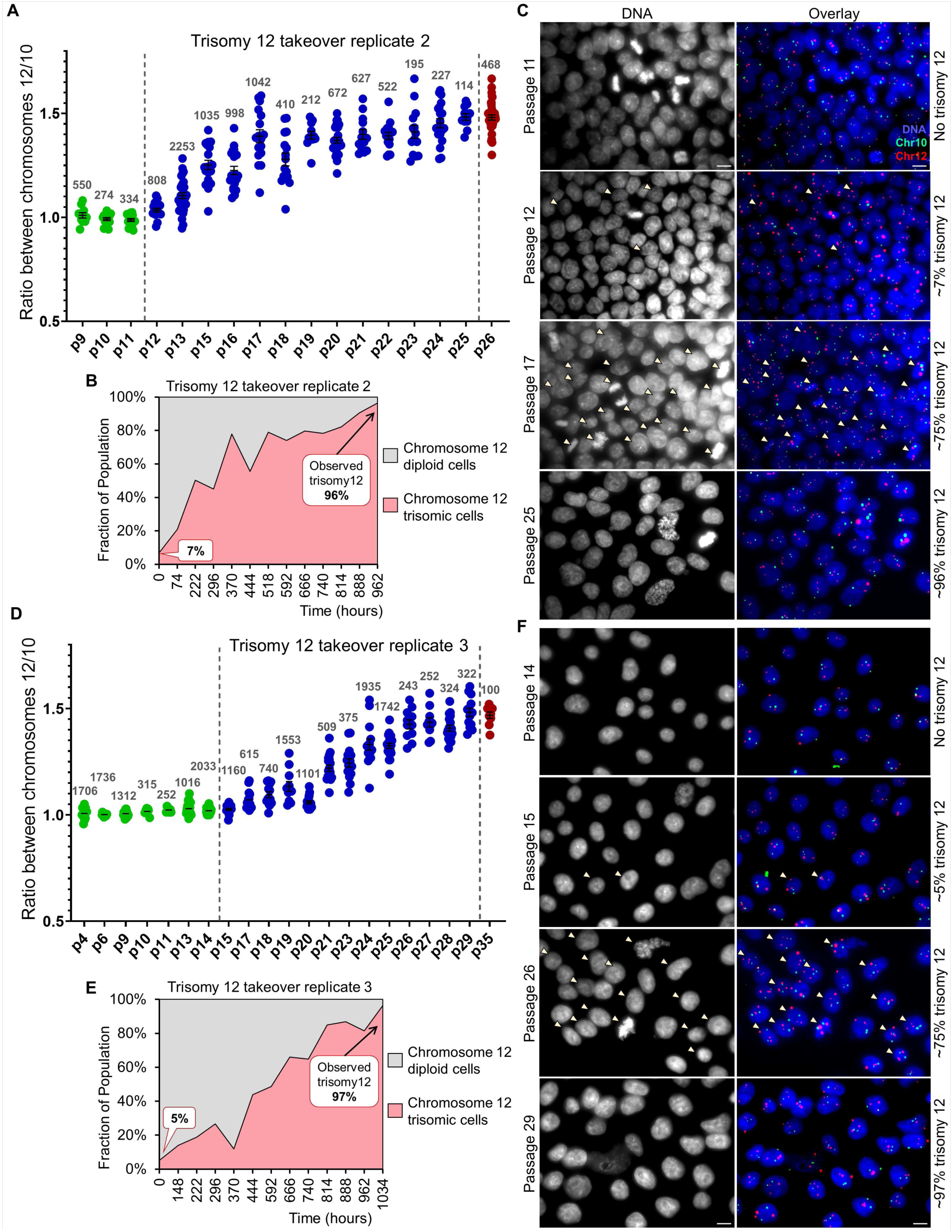
Chromosome 12 trisomic cells appear spontaneously in human iPSCs culture and become dominant within approximately 13 passages. **(A)** The observed trisomy 12 takeover during transition passages 12 to 25 in replicate 2 of the trisomy 12 takeover analysis. **(B)** The observed change in the ratio of chromosome 12 diploid and trisomic population over the course of 13 passages in replicate 2 of the trisomy 12 takeover analysis. **(C)** Representative images of cells from passages 11, 12, 17, and 25 in replicate 2 of trisomy 12 takeover analysis. Trisomy 12 is undetected in passage 11, but in transition passages 12 and 17, we observed approximately 7% and 75% of trisomy 12, respectively. Trisomy 12 then dominated (∼96%) by passage 25. **(D)** The observed trisomy 12 takeover during transition passages 15 to 29 in replicate 3 of the trisomy 12 takeover analysis. In **(A)** and **(D)** each dot on the graph represents a microscope field of analyzed cells, with the numbers above indicating the total number of analyzed cells. Error bars represent SEM. **(E)** The observed change in the ratio of chromosome 12 diploid and trisomic cells over the course of 14 passages in replicate 3 of the trisomy 12 takeover analysis. **(F)** Representative images of cells from passages 14, 15, 26, and 29 in replicate 3 of the trisomy 12 takeover analysis. In transition passages 15, and 26 we observe respectively ∼5% and ∼75% of trisomy 12. Trisomy 12 gained dominance (∼97%) at the passage 29. In **(C)** and **(F)**, DNA labeling is shown in grey and the overlays merge DNA (blue), chromosome 12 centromeres (red), and chromosome 10 centromeres (green). Arrows on the overlays indicate chromosome 12 trisomic cells. Images for ratio calculations were taken using a Zeiss Axioplan II microscope with 63× and 100× objectives and a Nikon TiE microscope using 40× or 63× objectives. Scale bar = 10 µm. Passage numbering reflects the number of passages performed during the course of experiments starting from passage 3 of human iPS cell line AICS-0012.

**Fig. S4.**
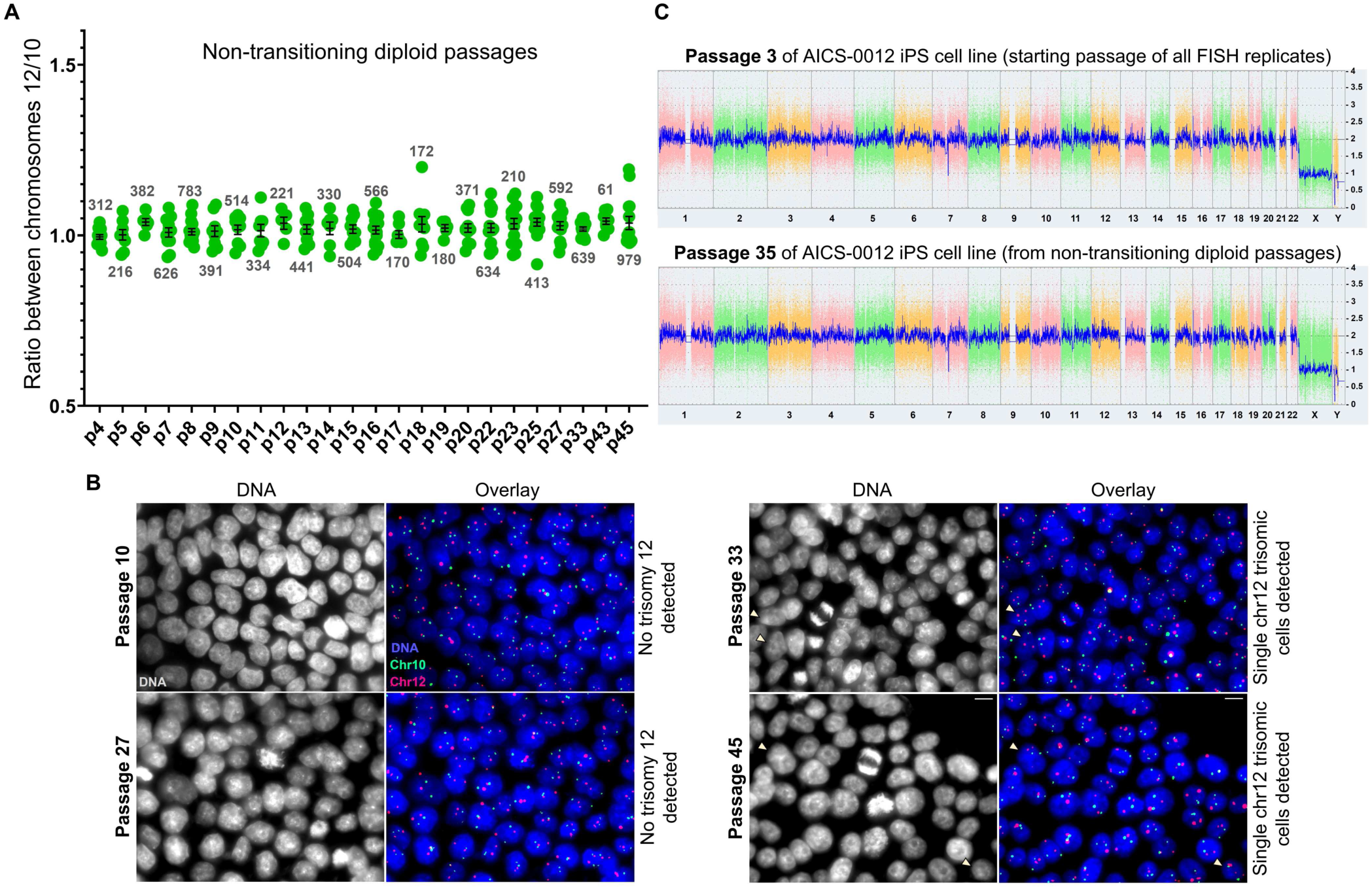
Chromosome 12 trisomic cells did not overtake the culture only one out of eight times, demonstrating that trisomy 12 takeover is the most frequent outcome. **(A)** The only passaging course where trisomy 12 takeover is undetected until at least passage 45. Each dot on the graph represents a microscope field of analyzed cells, with the numbers above indicating the total number of analyzed cells. Error bars represent SEM. **(B)** Representative images of cells from passages 10, 27, and passages 33 and 42, where only single chromosome 12 trisomic cells were detected. DNA labeling is shown in grey and the overlays merge DNA (blue), chromosome 12 centromeres (red), and chromosome 10 centromeres (green). Arrows on the overlays indicate chromosome 12 trisomic cells. Images for ratio calculations were taken using a Zeiss Axioplan II microscope with 63× and 100× objectives and a Nikon TiE microscope using 40× or 63× objectives. Scale bar = 10 µm. **(C)** The whole genome view of passage 3 and passage 35 after KaryoStat+ analysis performed on the non-transitioning diploid cell population. The whole genome view displays all somatic and sex chromosomes. The signal plot (right y-axis) is the smoothing of the log2 ratios which depict the signal intensities of probes on the microarray. A value of 2 represents a normal copy number state. The pink, green, and yellow colors indicate the raw signal for each individual chromosome probe, while the blue signal represents the normalized probe signal. Due to a lower probe density on the telomere ends and centromeres, the resolution in those locations on some chromosomes is lower. The deletion in chr Y is caused by the derivation of the AICS-0012 cell line from the WT cell line, which is known to have this genotype. No additional chromosomal gains or losses were detected in passages 3 and 35. Passage numbering reflects the number of passages performed during the course of the experiment starting from passage 3 of human iPS cell line AICS-0012.

**Fig. S5.**
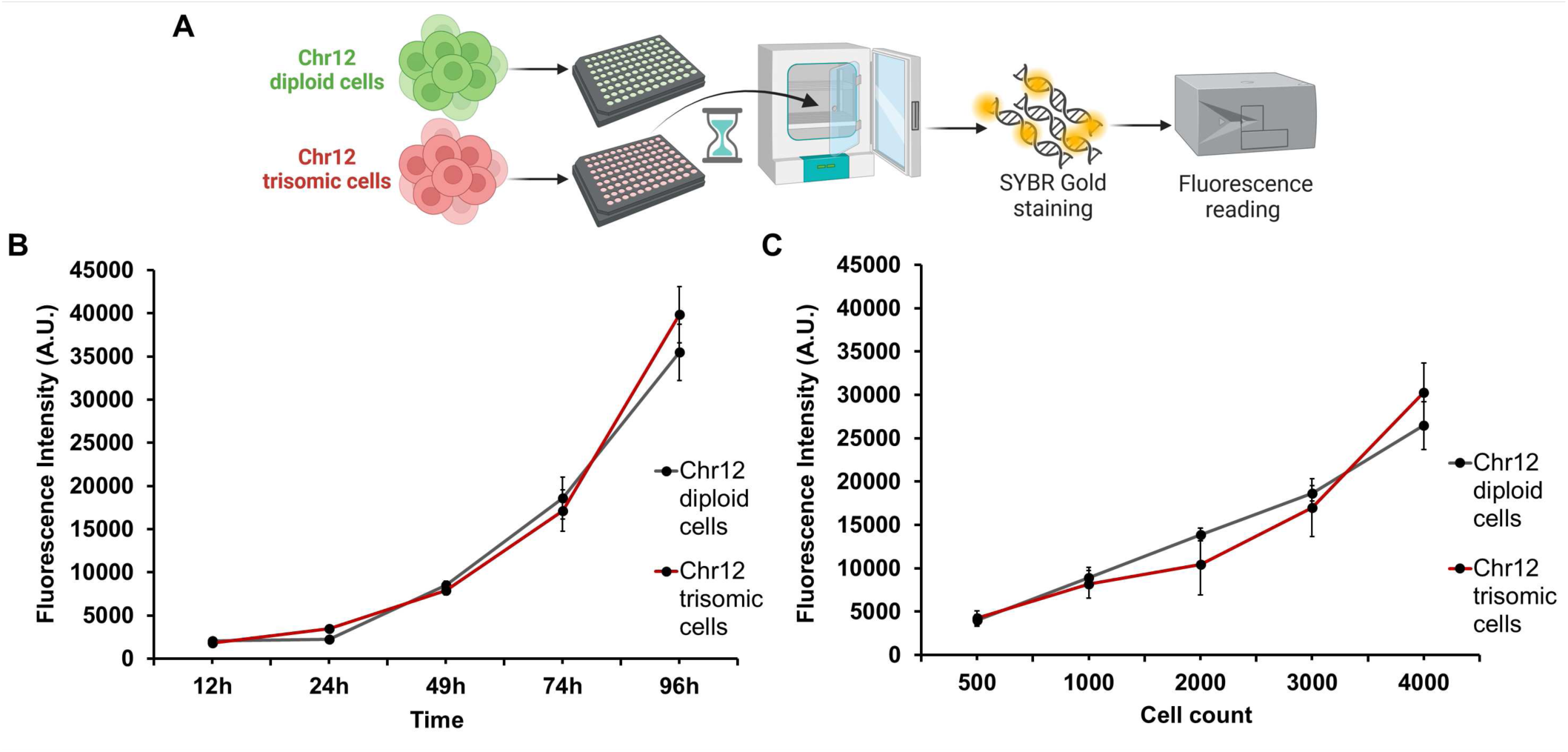
**(A)** Schematic of fluorescent DNA labeling using SYBR Gold to quantify the DNA content in diploid and trisomic iPS cell populations. **(B)** and **(C)** Two types of microwell growth assays conducted on diploid (grey line) and trisomic passages (red line). Initial seeding in **(B)** was 1700 cells per microwell, and cell counts were measured by fluorescence intensity at indicated time points on the x-axis. In **(C)**, varying cell amounts were seeded per microwell for the same growth time (70h). Doubling times were calculated using both growth assay types.

**Fig. S6.**
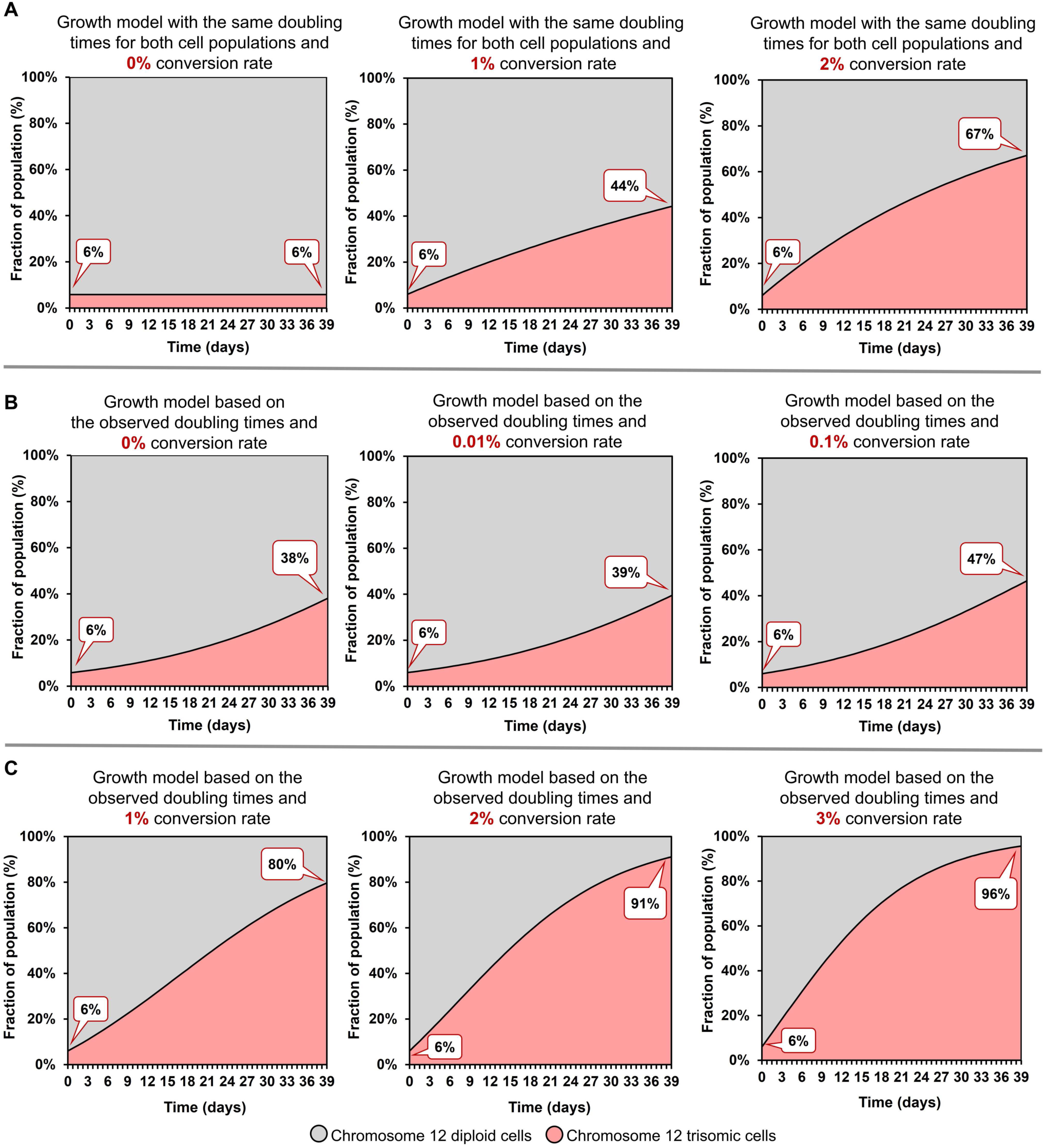
Mathematical modeling highlights the combined contributions of growth rate differences and high frequency of mis-segregation events in trisomy 12 dominance. **(A)** Simulated outgrowth of trisomic cells over diploid in the absence of a doubling time difference using conversion rates of 0%, 1%, and 2% per cell division. Even at a 2% conversion rate, trisomic cells fail to reach 70% of the population within 13 passages (39 days in cell culture), highlighting the necessity of faster growth rates of trisomic cells for trisomy 12 dominance during that time. **(B)** Simulated outgrowth of trisomic cells incorporating the observed doubling time difference between diploid and trisomic populations and conversion rates of 0%, 0.01%, and 0.1%. Trisomic cells do not reach 50% dominance within 13 passages (39 days in cell culture), demonstrating that a high frequency of chromosome 12 mis-segregation events is also essential for rapid trisomy 12 takeover. **(C)** Comprehensive mathematical models incorporating both, the doubling time difference and high conversion rates of diploid-to-trisomic cells (1%, 2%, and 3%). Only when both mechanisms are combined does the model replicate the observed rapid dominance of trisomic cells, reaching almost 100% dominance within 13 passages (39 days in cell culture).

